# Prophage-DB: A comprehensive database to explore diversity, distribution, and ecology of prophages

**DOI:** 10.1101/2024.07.11.603044

**Authors:** Etan Dieppa-Colón, Cody Martin, Karthik Anantharaman

## Abstract

**Background:** Viruses that infect prokaryotes (phages) constitute the most abundant group of biological agents, playing pivotal roles in microbial systems. They are known to impact microbial community dynamics, microbial ecology, and evolution. Efforts to document the diversity, host range, infection dynamics, and effects of bacteriophage infection on host cell metabolism are extremely underexplored. Phages are classified as virulent or temperate based on their life cycles. Temperate phages adopt the lysogenic mode of infection, where the genome integrates into the host cell genome forming a prophage. Prophages enable viral genome replication without host cell lysis, and often contribute novel and beneficial traits to the host genome. Current phage research predominantly focuses on lytic phages, leaving a significant gap in knowledge regarding prophages, including their biology, diversity, and ecological roles.

**Results:** Here we develop and describe Prophage-DB, a database of prophages, their proteins, and associated metadata that will serve as a resource for viral genomics and microbial ecology. To create the database, we identified and characterized prophages from genomes in three of the largest publicly available databases. We applied several state-of-the-art tools in our pipeline to annotate these viruses, cluster and taxonomically classify them, and detect their respective auxiliary metabolic genes. In total, we identify and characterize over 350,000 prophages and 35,000 auxiliary metabolic genes. Our prophage database is highly representative based on statistical results and contains prophages from a diverse set of archaeal and bacterial hosts which show a wide environmental distribution.

**Conclusion:** Prophages are particularly overlooked in viral ecology and merit increased attention due to their vital implications for microbiomes and their hosts. Here, we created Prophage-DB to advance our comprehension of prophages in microbiomes through a comprehensive characterization of prophages in publicly available genomes. We propose that Prophage-DB will serve as a valuable resource for advancing phage research, offering insights into viral taxonomy, host relationships, auxiliary metabolic genes, and environmental distribution.

## INTRODUCTION

Phages constitute the most abundant group of biological agents on the planet, with conservative estimates suggesting numbers of 10^30^ around the globe^1^. As consequence of their sheer abundance, they have significant impacts across all microbial systems, hence they are of utmost importance in the study of the microbial world.

Phage impacts are dependent on the mode of bacteriophage infection which can be lytic, lysogenic, or chronic^2^. While lytic infection consists of viral particle production and host lysis to release the viral particles, the lysogenic mode of infection, consists of phage integration into the host cell genome. This integration is known as a prophage and allows the bacteriophage genome to be replicated without causing the lysis of the host by repressing lytic functions^2,3^. Phages capable of lysogeny are known as temperate viruses and can switch between the lytic and lysogenic cycles. The switch in mode of infection from lysogeny to lytic is known as induction and can be caused by different external factors, these can include, antibiotics^4^, UV rays^5^, reactive oxygen species^6^, changes in temperature^7^, changes in pH^8^, bacterial metabolites and products of host physiology^9^, but can also occur spontaneously^10^. In addition, in polylysogeny, an event in which there are multiple prophages in a single host, prophages can encode noncanonical induction pathways to outcompete each other^11^. The switch between modes of infection leads to changes in microbial communities that extend beyond virus-host interactions and have broader implications. Prophages are commonly present within the genomes of bacteria and archaea, with some cases showing they can make up to 20% of the genome^12^. In addition, most bacteria, are polylysogens^11^ and a well-studied case is the *E. coli* O157:H7 strain Sakai which has 18 prophages^13^. However, typically, a prophage genome represents only about 1% of the host’s genome^14^. Finally, in the chronic mode of infection, viral particles are continuously produced without lysis of the host^2^.

Prophages can impact their respective microbial systems in different ways. They can manipulate the host’s gene expression and function, affecting the host’s cellular processes; they can also alter the host’s physiological functions or introduce new functions^2^. At a broader level, prophages impact the structure, function, ecology, and evolution of microbial systems^2,10,15,16,17^. Specifically, by lysing microbes in competition, prophages can prompt shifts in community dynamics^2^. In host-associated systems, such as the human gut microbiome, they have the potential to impact the host’s physiology and health^10,18^. For example, their bacterial hosts’ genomes and phenotypes can undergo changes which has the potential for the emergence of strains and diversification; this has implications in the hosts’ virulence and antibiotic resistance^19^ which in turn could affect the overall health of hosts such as humans.

Phages can contain auxiliary metabolic genes (AMGs) which are involved in numerous metabolic processes and can alter host metabolism. AMGs are host-derived genes normally acquired by the phage through recombination and which can be expressed during infection by the phage to improve its fitness^20^. Their presence is of high relevance given that these phage-encoded genes are involved in host metabolism and respond rapidly to environmental cues^21^. Additionally, through the incorporation of AMGs, prophages have the potential to influence ecosystem biogeochemistry, thereby impacting global biochemical processes. However, much remains to be unraveled to fully understand the extent of their impact^22^. Overall, our knowledge about prophages and their AMGs and the relationships between their host remains poor. Even though we know about certain relationships such as prophages being more frequent in pathogenic and fast growing bacteria^23^, or that viral lifestyle is a major driver of AMG composition^24^, we still need to strive for a more universal understanding regarding of such relationships.

Current limitations in prophage identification arise from the fact that viruses are polyphyletic, meaning they have multiple evolutionary ancestors. As a consequence, they do not display universal markers that ease their identification in datasets and hence, the use of databases only allows for the identification of a few. As a result, the ability to discover novel viruses remains restrained. Metagenomic studies have provided extensive insights into viruses, giving rise to the concept of ‘viral dark matter.’ This term refers to the unknown identities and functions of viruses and their proteins, the majority of which have no known function^12,25^. Chevallereau et al. defined the concept of ‘viral dark matter’ as viral species that have not been characterized but for which their existence has been revealed by metagenomic sequencing, or phage genes that have no assigned functions^3^. Surprisingly, in certain datasets, ‘viral dark matter’ may constitute up to 90% of sequences. Studies have identified three factors contributing to the existence of ‘viral dark matter’: the divergence and length of virus sequences, limitations in alignment-based classification, and the inadequate representation of viruses in reference databases^26^. As a result, a considerable number of viruses resist taxonomic classification or association with a bacterial or archaeal host which underscores the necessity to expand our knowledge to fully comprehend their potential effects on their respective microbial systems.

Several methods have been developed to circumvent the limitations of the exclusive use of viral hallmark genes and homology-based methods, and many integrate a combined approach to complement sequence similarity comparisons. Notable examples include the use of machine learning or the search for composition patterns such as k-mer frequency among others. Examples of tools that use a combined approach include DEPhT^14^, geNomad^27^, PHASTEST^28^, PhiSpy^19^, VIBRANT^20^, VirSorter2^29^, and VirFinder^30^. Although the methods used by these tools have been effective in identifying phages, much remains undiscovered. This can be facilitated by the creation of standardized databases that can drive the discovery of novel phage signatures.

Despite the numerous methods/tools that have been developed for the study of phages, the number of databases lags behind. For the most part, current databases are associated with software and their purpose is to be used as a reference for homology-based comparisons. Only a small subset of databases exist as catalogues that can be used to explore the phages contained within. And from this subset, only a few are comprehensive as several databases focus on a single microbial system, such as the gut microbiome or marine environments. Examples of comprehensive prophage studies/datasets focused on specific microbial systems include Marine Temperate Viral Genome Dataset (MTVGD)^31^ and Human Gut-derived Bacterial Prophages^32^. Examples of comprehensive databases that contain prophage sequences are Microbe Versus Phage database^33^ intended for exploring the relationships between phages and their hosts; IMG/VR v4^34^, which contains a significant number of prophages and metadata such as environment; and PhageScope^35^, a bacteriophage database encompassing temperate and lytic viruses.

Here, we present Prophage-DB, a comprehensive database of prophages that will serve as a standardized resource to facilitate viral diversity and ecology studies.

## METHODS

### Data collection

To create Prophage-DB, we identified prophages from publicly available prokaryotic genomes from three databases: Genome Taxonomy Database (GTDB)^36^ (release 207), National Center for Biotechnology Information (NCBI) Reference Sequence (RefSeq) database^37^ (accessed March 2023), and Searchable Planetary-scale mIcrobiome REsource (SPIRE)^38^. The prokaryotic genomes contained both archaeal and bacterial representative genomes and were used as the initial input. To generate the dataset, we applied several tools to identify and annotate viruses, cluster them, assign taxonomy and measure their quality (**Figure 1**). The SPIRE database also included a portion of genomes with no domain classification labeled as unknown (these are referred to as unclassified).

**Figure 1.**
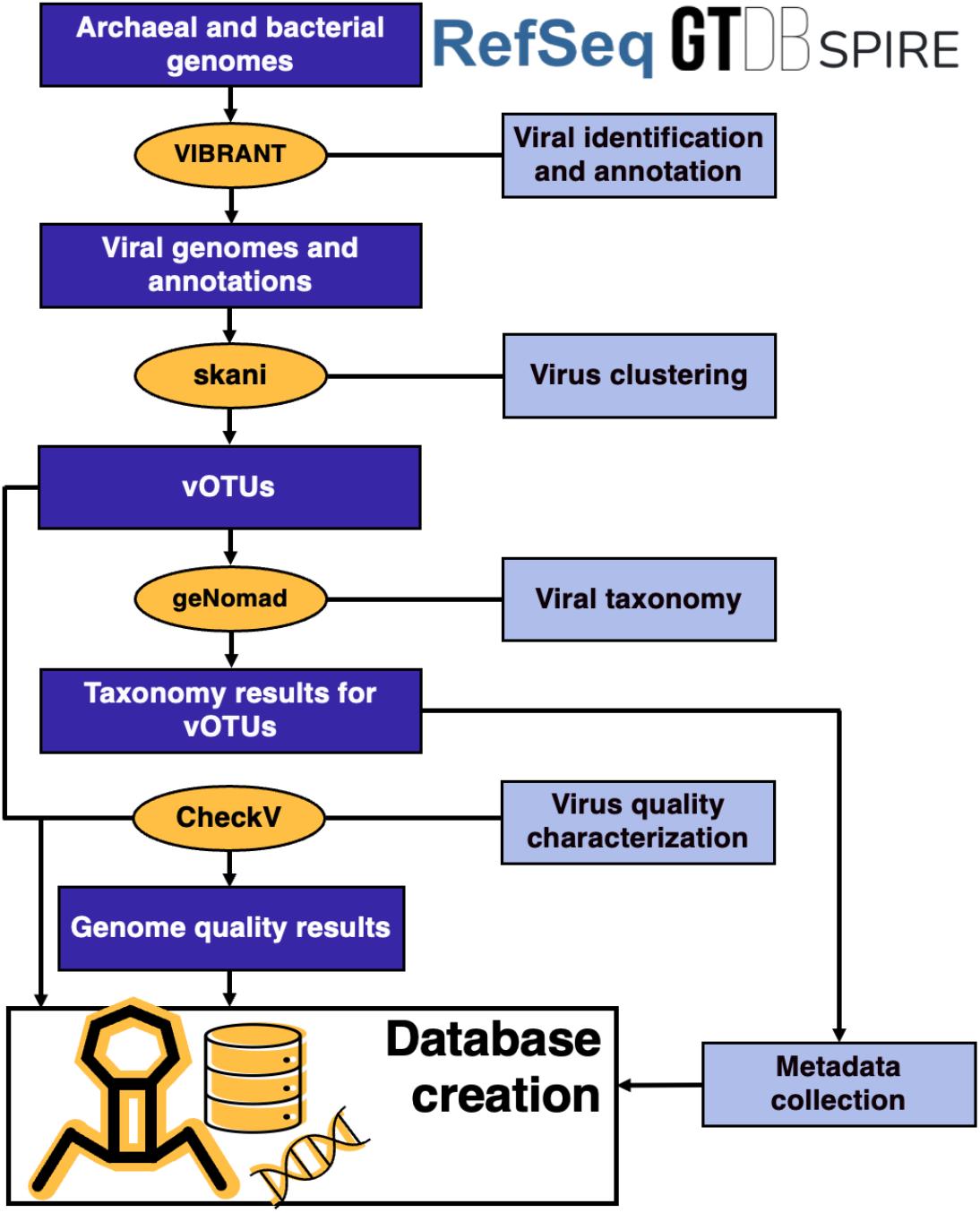
Visual representation of steps and software utilized to create database.

### Prophage identification

Prophage identification was carried out using VIBRANT (v1.2.1) which identifies and annotates viruses from nucleotide scaffolds. VIBRANT is able to detect both lytic and lysogenic viruses, however, given that all viruses were identified from prokaryotic genomes they were deemed as prophages. We used the default arguments when using VIBRANT (minimum scaffold length requirement = 1000 base pairs, minimum number of open readings frames (ORFs, or proteins) per scaffold requirement = 4).

### Virus clustering

Due to redundancy among the genomes in the three databases our next step was to cluster the identified viruses (VIBRANT output nucleotide files for phages). For this we used the algorithm and software, skani (v0.2.1)^39^, this tool performs average nucleotide identity (ANI) calculation using a k-mer scheme. We performed all-to-all comparisons using the skani default arguments, with exception to the alignment fraction argument which was set to 85 (--min-af 85). After obtaining ANI and alignment fraction, we removed viral sequences for which ANI was 100 and both the query and subject had at least 85 alignment fractions. In the removal of identical sequences, we prioritized SPIRE sequences over GTDB and NCBI sequences given that the bulk of the data originated from SPIRE.

### Viral taxonomy and metadata

After performing clustering to remove highly similar sequences, we utilized geNomad (v1.7.0)^27^ to perform taxonomic assignment of the viral genomes (specifically, we used the annotate module). In addition, we performed virus quality characterization using CheckV (v1.0.1)^40^ obtaining viral quality, completeness, and contamination. The workflow culminated with collecting metadata from GTDB and SPIRE which provided important information such as host taxonomy, host isolation source, and geographical coordinates. Additional metadata includes AMGs highlighted by VIBRANT, and metrics provided by CheckV (**Table S1, S2**).

### Representation analysis

We utilized the taxonomic assignment counts of the provirus subset of IMG/VR v4 (all sequences) and prophages in Prophage-DB. Given that IMG/VR is a flagship database in viral ecology and contains the largest uncultivated virus genomes (UViGs) collection^34^, we assume that it should be the closest database to being representative of real phage diversity. To determine representation, we followed the approach carried out by Titley et al.^41^ in which groups above the 1:1 line were classified as over-represented and those below as under-represented. We compared the logarithmic proportions of the prophages in our database to those in IMG/VR v4 across various viral taxonomic ranks. By comparing it to our observed counts, it allowed us to determine which groups were over and under-represented.

### Statistical analysis

We employed random sampling with replacement (bootstrap value of 10,000) on the taxonomic assignments of the provirus subset from IMG/VR v4 and prophages in Prophage-DB. This step was executed using a custom Python script, where the sampling was performed utilizing the .sample() method from the Pandas library based on the taxonomic counts for each taxonomic rank. Considering the comprehensiveness of IMG/VR v4, we compared its results with those from our database to validate our observations.

### AMG filtering

AMGs were identified with VIBRANT. For filtering AMGs, we followed the steps outlined in Zhou et al^42^: 1)AMGs at either end of the scaffold were filtered out; 2) AMGs with VIBRANT KEGG^43^ or Pfam^44^ v-scores equal or greater than 1 were filtered out; 3) AMGs that had four genes (upstream or downstream) with KEGG v-score smaller than 0.25 were filtered out.

## RESULTS

### Prophage identification

After viral identification and clustering, we obtained 356,776 prophage sequences, most of which originated from bacterial hosts (323,608 prophages from bacterial hosts). The second largest group consisted of prophages from unclassified hosts (21,226 prophages), while prophages in archaeal hosts (11,942) comprised a smaller subset (**Figure 2A**) (**Table S1**). Across all taxonomic ranks, except species, the majority of prophages in our database were associated with a host with known taxonomy (**Figure 2B**). In comparison to the reported taxonomic ranks in GTDB, we found that at least 50% of them contained prophages to the family level for both archaea and bacteria (**Table S3**). Phage taxonomy was assigned most down to the class level, while at the order and family level, only a small subset of phages had unassigned taxonomy (**Figure 2C-E**). Next, we used CheckV to assess the quality of our prophage genomes. Most genomes were labeled as low quality, and with a low completeness even while they were detected in complete prokaryotic genomes (**Table S4**). These results serve as evidence of our poor knowledge of prophages and viral dark matter and highlight the need for more in-depth studies that search for novel signatures associated with prophages.

**Figure 2.**
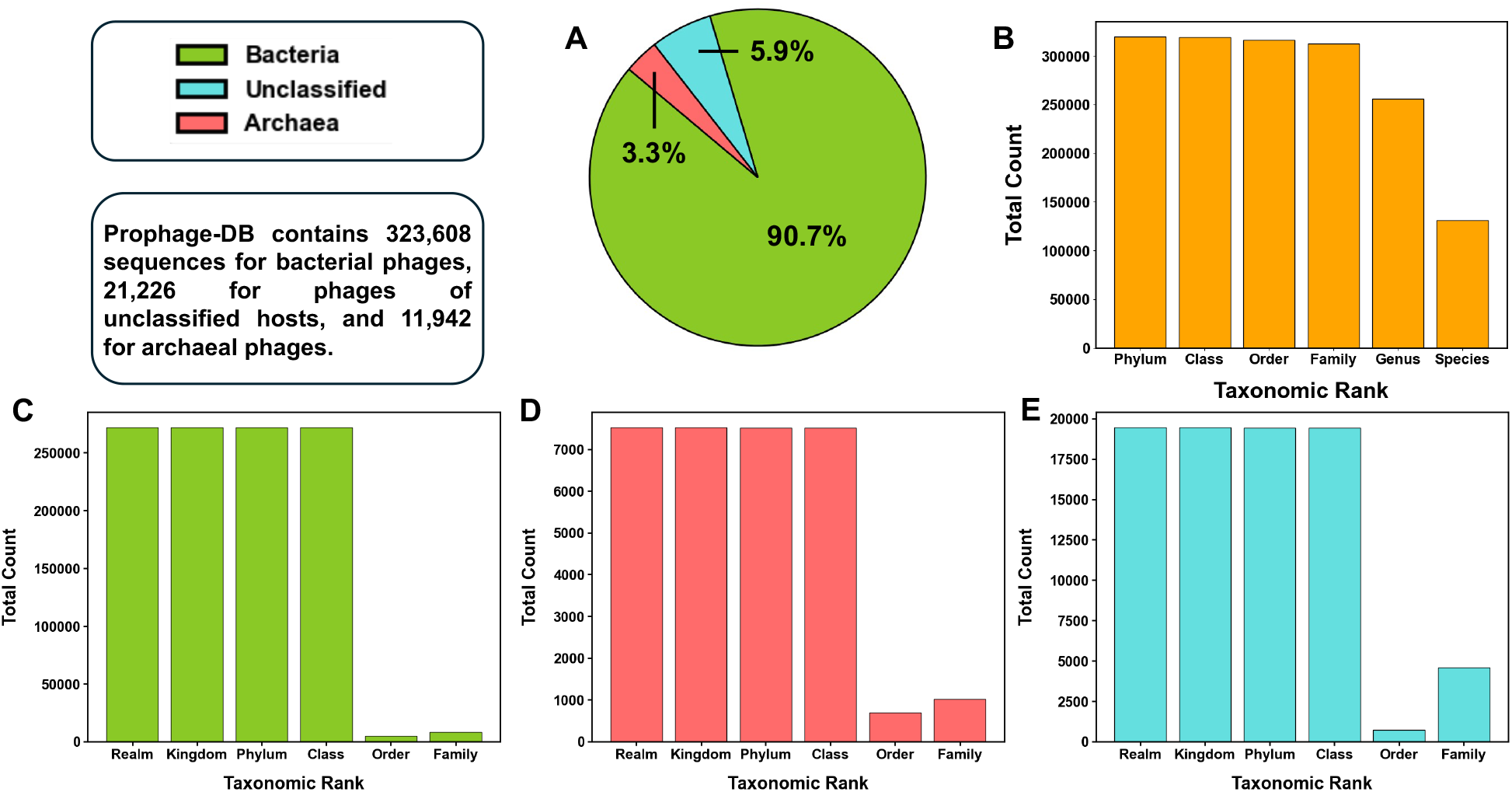
Prophage-DB general metrics. A) Percentages of the three host groups in the database (archaea, bacteria, and unclassified hosts). B) Total count of prophage hosts with available metadata regarding a specific host taxonomic rank. C) Counts of taxonomic ranks for which prophages have been assigned to in the bacterial host group. D) Counts of taxonomic ranks for which prophages have been assigned to in the archaeal host group. E) Counts of taxonomic ranks for which prophages have been assigned to in the unclassified host group.

### Prophage taxonomy in Prophage-DB

A significant proportion of the prophage sequences (84%) in our database were assigned to a taxonomic rank. At higher taxonomic ranks, phage taxonomy followed a similar distribution across archaeal and bacterial groups (**Figure S1**). In archaeal hosts, the Duplodnaviria (realm), Heunggongvirae (kingdom), Uroviricota (phylum), and Caudoviricetes (class) constitute around 92% of the observations. Similarly, in bacterial hosts, the same groups constitute around 98% of the observations. The same pattern is also observed in prophages found in unclassified hosts. This imbalance is evidence of the bias towards studying certain groups and reveals groups of interest that show promise in the discovery of novel prophage signatures. In contrast, at the order and family level, the taxonomic ranks show a more balanced distribution in which most phages were not in a single group but in several. This was observed for prophages in archaea, bacteria, and unclassified hosts (**Figure 3**). At the order level, most prophages in bacteria and unclassified hosts belong to Crassvirales. In contrast in archaeal phages, the group Haloruvirales were the most abundant (**Figure 3**). Interestingly, the group Haloruvirales has little representation in bacteria. Similarly, the order Tubulavirales was frequent in bacteria but had low frequency in archaea. When compared to archaeal phages, prophages in bacteria showed more diversity (there were 7 groups found only in bacteria, highlighted with color). At the family level, we observed more differences across the three groups (**Figure 3**). 17 groups were found only in bacterial phages, 2 (Globuloviridae and Malacoherpesviridae) only in archaeal phages, and one (Retroviridae) only in phages from unclassified hosts. Inoviridae, Pleolipoviridae, and Kyanoviridae were the families with most counts for bacteria, archaea, and phages from unclassified hosts, respectively. Just as we observed at the order level, some groups predominant in bacteria have low frequencies in archaea and vice versa. This was the case for Pleolipoviridae which is predominant in archaeal hosts but not in bacteria. On the other hand, Inoviridae was the group with most counts in bacterial hosts but had very low frequency in archaeal hosts.

**Figure 3.**
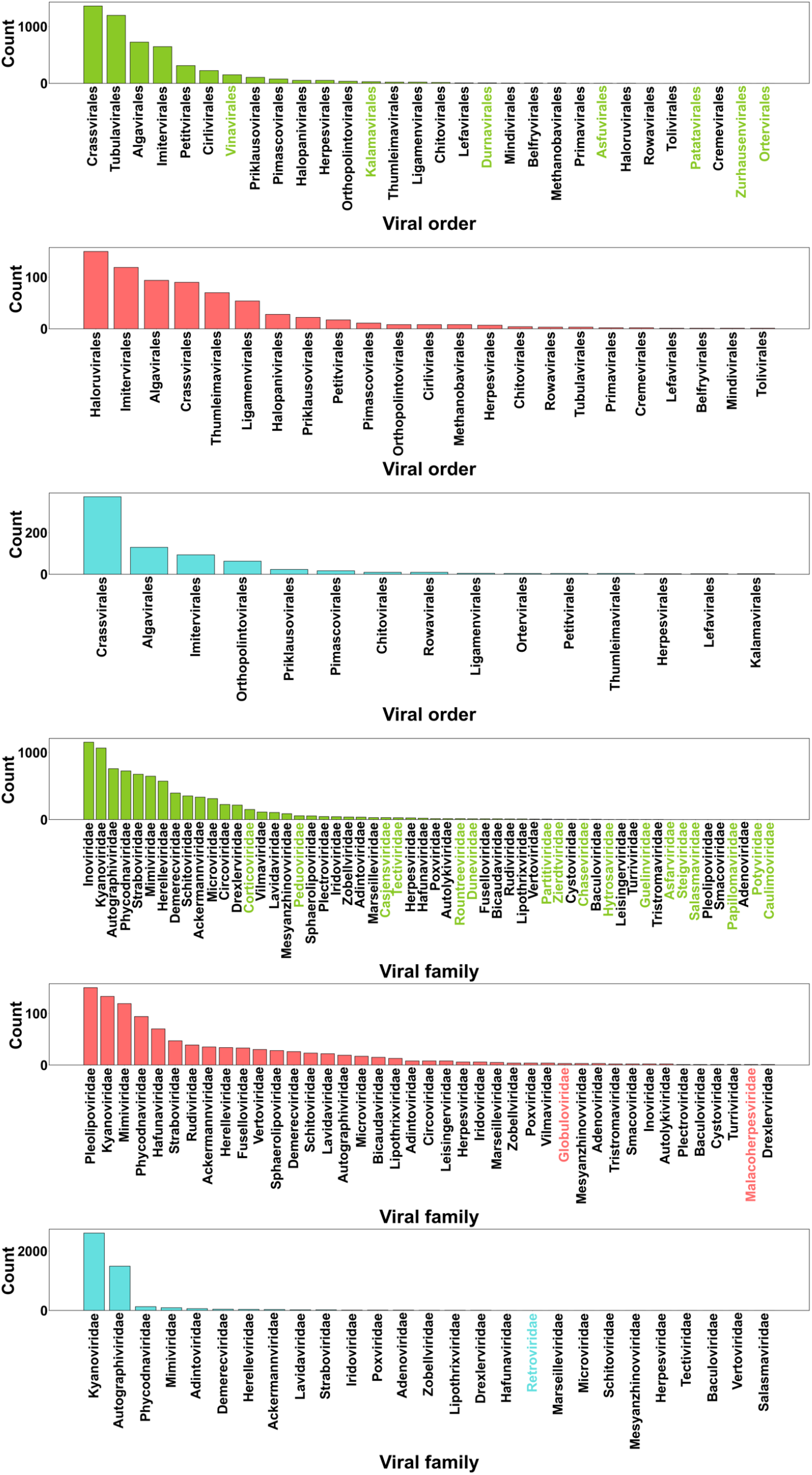
Order and family counts for prophage groups. Words in color represent orders found only in one group. Y-axis represents the counts of each taxonomic group; x-axis contains the names of taxonomic groups. Green, pink, and blue colors correspond to bacterial, archaeal, and phages from unclassified hosts, respectively.

### Environmental distribution of prophages in Prophage-DB

Next, we analyzed the prophage host distribution. Our database contained 149,796 hosts which were widely distributed throughout the world (**Figure 4A**) (**Table S1**). However, it was noticeable that developed countries contained more samples. To be more specific we separated our data by environment or isolation source (**Figure 4B**). We focused on 5 main environments of interest: host-associated, marine, fresh-water, terrestrial, and anthropogenic. As expected, prophages of bacteria were mostly related to host-associated bacteria (40%). Meanwhile, in archaeal hosts, more than half of the prophages (57%) were associated with aquatic environments. Similarly, in unclassified hosts, most prophages were associated with aquatic environments, and specifically, 75% came from marine environments. The anthropogenic isolation source was the smallest group for all three groups.

**Figure 4.**
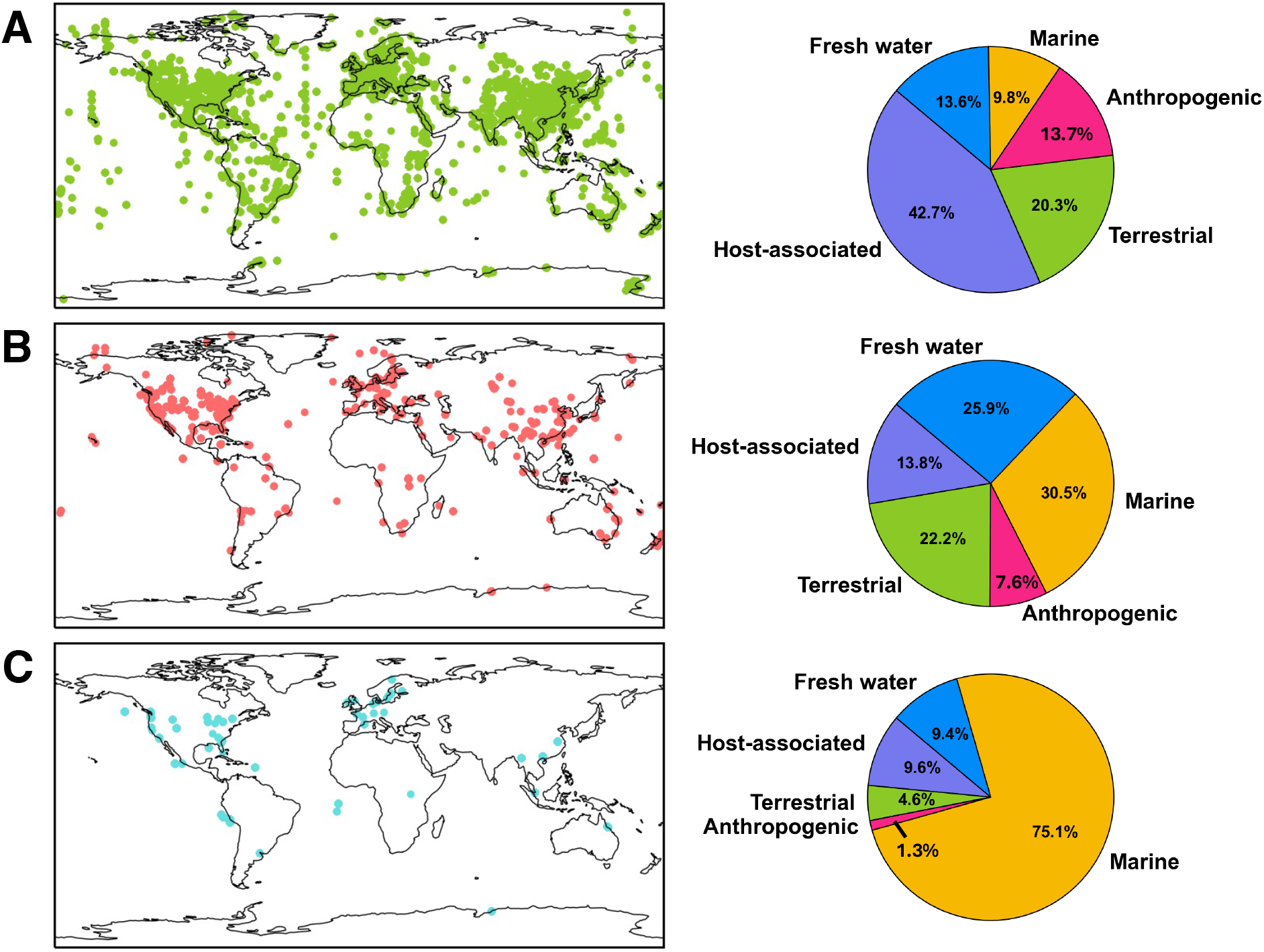
Phage-host distribution and isolation source. A) Locations of bacterial hosts’ isolation sources (left panel). Percentages of selected environments for bacterial hosts (right panel). B) Locations of archaeal hosts’ isolation sources (left panel). Percentages of selected environments for archaeal hosts’ (right panel). C) Locations of unclassified hosts’ isolation sources (left panel). Percentages of selected environments for unclassified hosts (right panel).

### Prophage count analysis in Prophage-DB

Next, we evaluated prophage distribution across host taxonomic ranks, i.e. the number of prophages per host. In these analyses, we utilized a non-dereplicated dataset, as using the prophages directly from our database would not accurately reflect prophage distribution. Specifically, we looked at archaeal and bacterial phyla, and accompanied this data with the count of the hosts (white bars) (**Figure 5**)). In addition, we extended this analysis to other host taxonomic ranks (**Table S5, Table S6, Figure S2-S4**). In archaea, the most common host phylum, Thermoproteota, had the largest prophage count (**Figure 5B**). This was also observed for Pseudomonadota, the most common phylum in bacteria in our database (**Figure 5A**). For the most part, all other phyla followed this pattern, more hosts meant a greater number of prophages.

**Figure 5.**
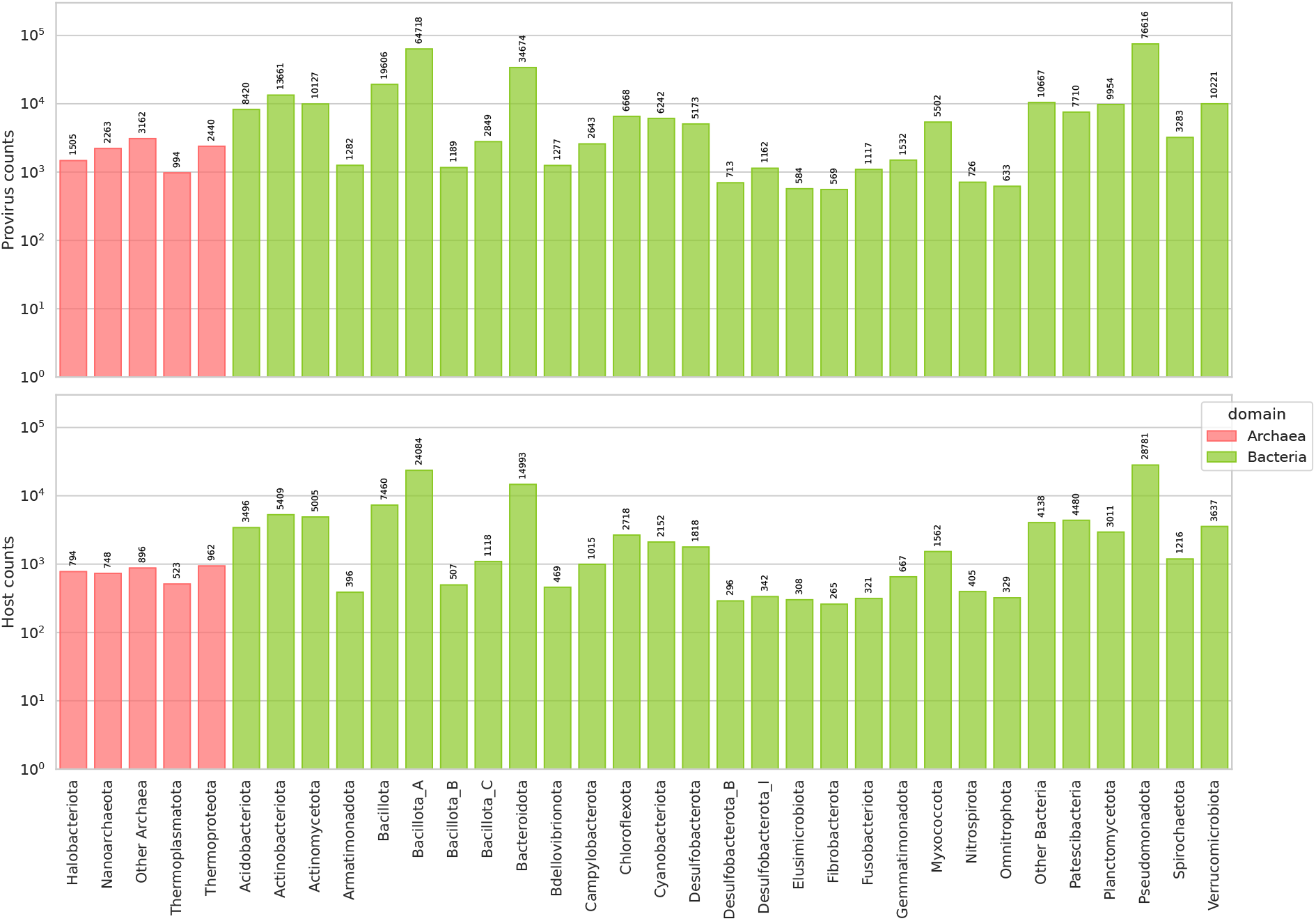
Prophage distribution across host taxa. A) Host and prophage count in bacteria and archaea. Only phyla with the greatest number of prophages are shown.

However, some phyla showed higher ratios by having a high number of prophages. The group with the highest prophage ratio was Asgardarchaeaota in archaea. Numerous phyla had low prophage counts in bacteria.

### Representation analyses of genomes in Prophage-DB

Next, we analyzed the representation status of viral groups. To determine representation, the baseline is that groups above the 1:1 line are classified as over-represented and those below as under-represented. We compared the logarithmic proportions of the prophages in our database to the provirus subset in the Integrated Microbial Genomes Viral Resources v4 (IMG/VR) across various viral taxonomic ranks (**Figure 6**). At the class, order, and family levels, archaeal phages tended to be over-represented, on the other hand, at these levels, bacterial phages tended to be under-represented. This analysis was also carried out by separating the viral taxonomic ranks by environment (**Table S7-S12, Figure S5-S14**).

**Figure 6.**
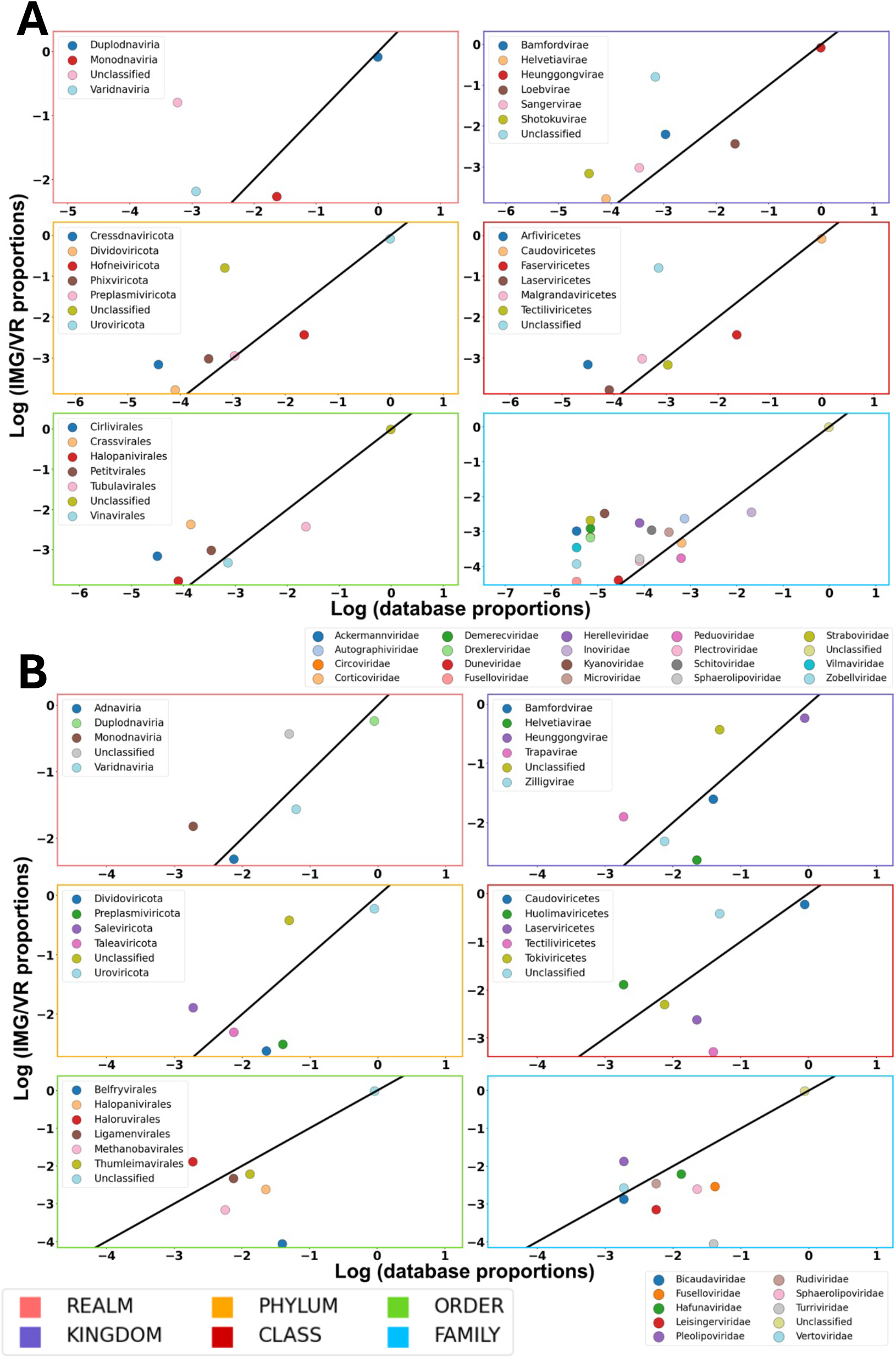
Representation analysis comparing proportion of taxonomic groups in our database to the proportions in IMG/VR database. Axes contain log transformed values of the counts. Values above the 1:1 line are considered over-represented, those below, are under-represented. Panel A shows results for bacterial phages; panel B shows results for archaeal phages. Margin colors represent taxonomic levels.

### Auxiliary Metabolic Genes

We identified a total of 35,990 AMGs. 33,617 were found in bacterial phages, 1,327 in archaeal phages and 1,045 in phages of unclassified hosts (**Table S2**). We analyzed how AMGs are distributed through five selected environments (host-associated, marine, fresh-water, terrestrial, and anthropogenic) (**Figure 7A**). Additionally, we analyzed AMG groups and their distribution across archaeal and bacterial hosts (**Figure 7B**). AMG distribution was mostly similar across all environments. In addition, AMG distribution by host was mostly similar across phyla.

**Figure 7.**
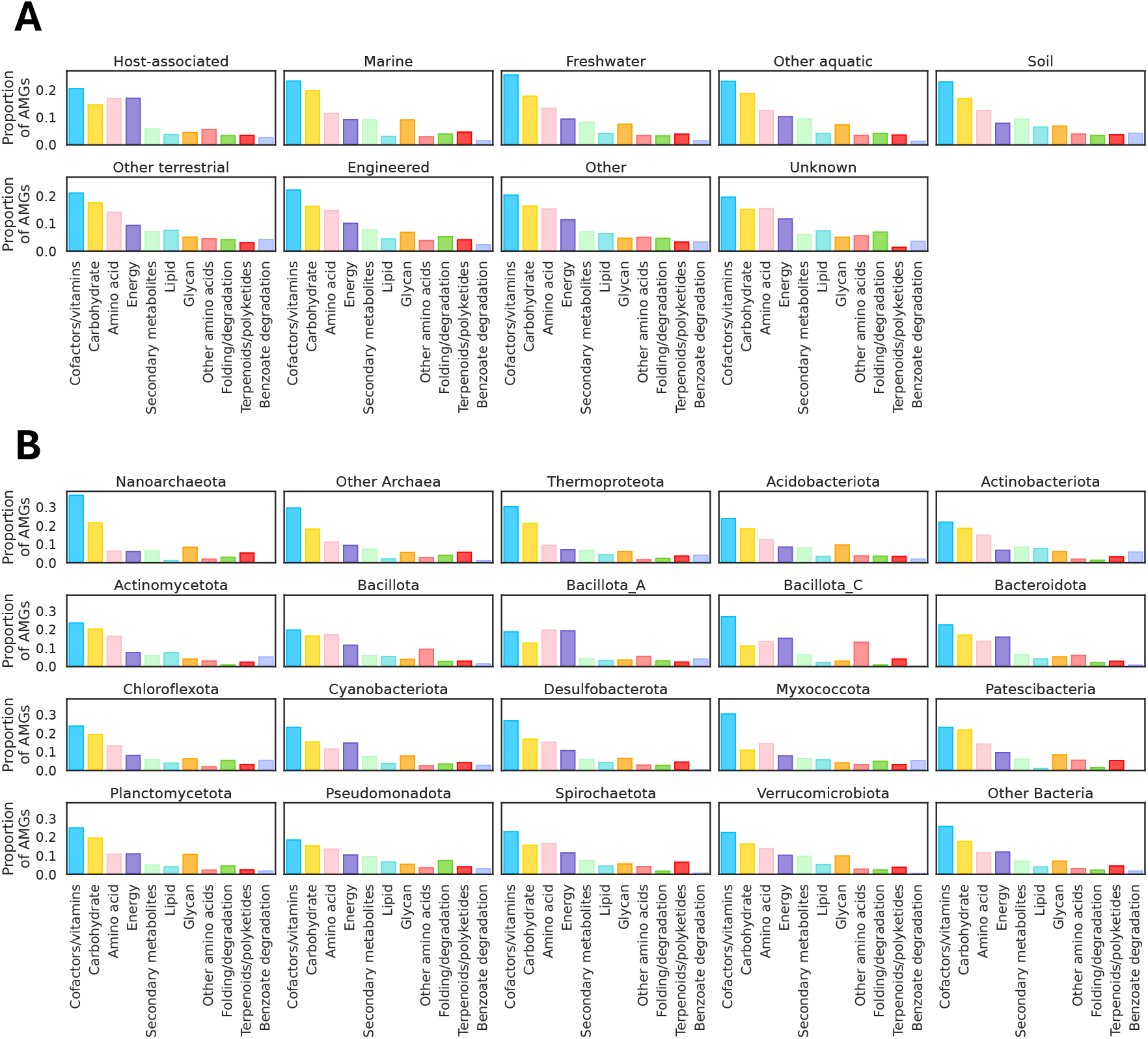
AMG distribution across environments and host taxa. A) AMG distribution across different environments. The group *Other* represents data that did not fall into any of the shown categories. The *Unknown* category represents data for which environment is labeled as unknown. B) AMGs distribution by host phyla (data shown is only for phyla with 250 or more counts). Nucleotide metabolism AMGs were not considered in this analysis.

### Statistical analysis of observed virus counts in Prophage-DB

We utilized random sampling with replacement (bootstrapping value of 10,000) on taxonomic counts belonging to our database and IMG/VR. Given that IMG/VR is a robust dataset it served as a baseline to identify expected counts in our database. In other words, we expected to observe similar counts between IMG/VR and our database. At the family level, both expected and observed counts are similar and the majority fall into the same group (in this case most phages were unclassified). This trend is observed at the rest of the taxonomic levels (**Figure 8, Tables S13-S24, Figures S15-S24**). We carried out this same analysis but separated the data into the 5 previously shown environments. We observed the same trend for the majority but expected and observed counts tended to deviate more than when analyzing all counts.

**Figure 8.**
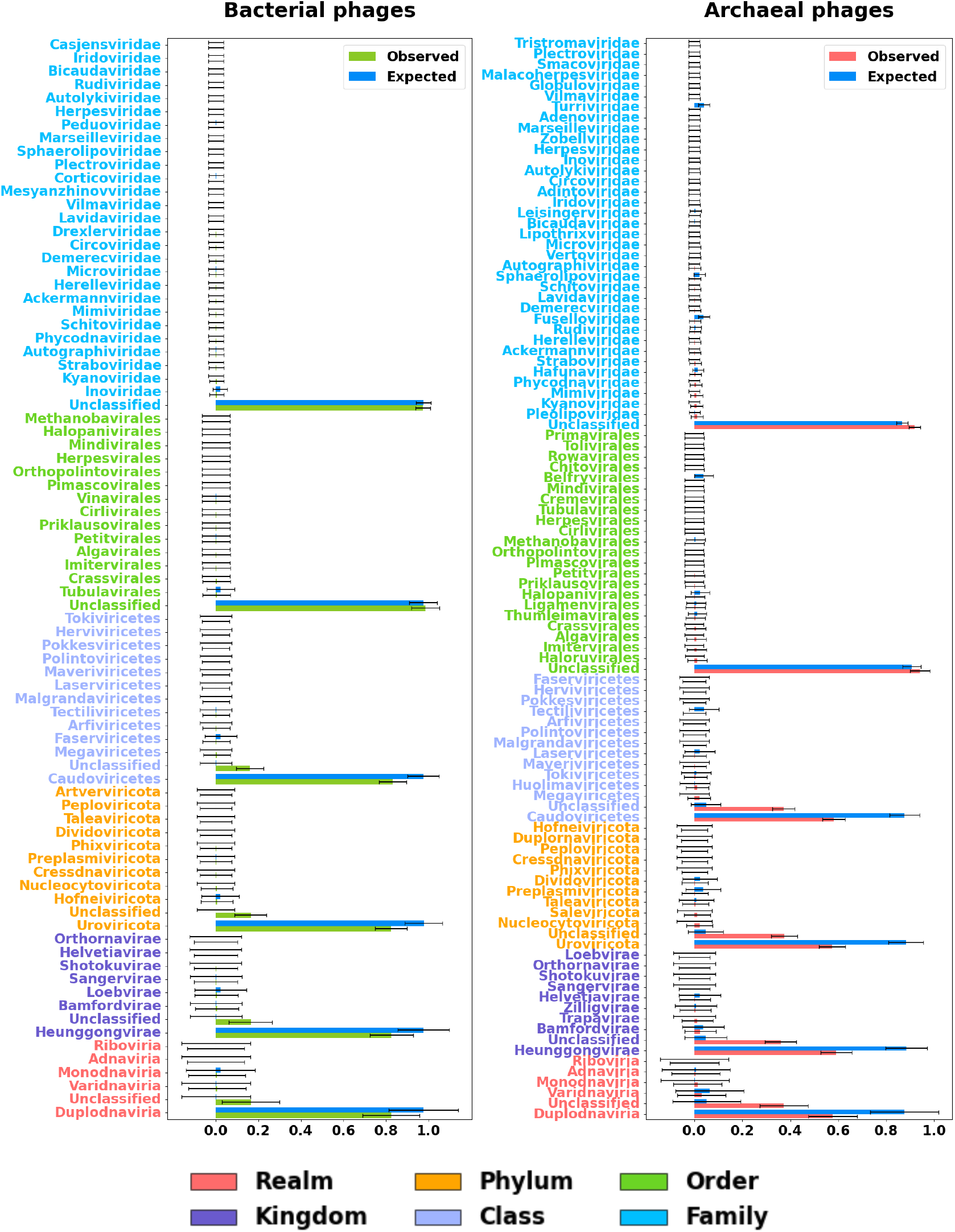
Comparison between IMG/VR counts (expected) and Prophage-DB counts (observed). X-axis represents the percentage. Random sampling with replacement was used (Bootstrap value of 10,000). Error bars represent the standard error. Y-axis colors represent taxonomic rank (shown in legend).

## DISCUSSION

As expected, a great portion of prophages originated from bacterial hosts while only a small percentage originated from archaeal hosts as the databases used in this study mainly contained bacterial genomes (**Table S1**). The small number of archaeal prophages in our database could stem from the historical focus on bacteria. Given that archaeal viruses are estimated to be as abundant as their bacterial counterparts^45^ we should expect to observe a similar amount unless they are inherently less diverse. The focus on bacteria has resulted in a gap in our knowledge and classification systems for prophages in the archaeal domain. A likely contributing factor are that techniques used to study phage in experimental settings are biased towards bacteria-infecting phages^25^. The small percentage of archaea in our database highlights the importance of further research in order to characterize more archaea and detect more archaeal prophages.

Regarding virus taxonomy, we observed a bias towards double-stranded viruses, resulting in most viruses being classified into a single realm: Duplodnaviria (all double-stranded DNA viruses). This result was not surprising as tailed phages with double-stranded DNA genomes are the most prevalent group of phages in public databases^1^. A likely factor contributing to the high prevalence of tailed phages with double-stranded DNA genomes could be a result of biased virology techniques and approaches. For example, in soil environments there is a bias towards this group as microscopic counting methods primarily detect tailed and encapsulated phages^25^. Ultimately, the majority of viral signatures used in viral identification also belong to this group. As expected, most of the identified prophages are in the Heunggongvirae kingdom (viruses containing the HK97 fold major capsid protein) given that it’s the only kingdom in the Duplodnaviria realm.

Despite these limitations, throughout our analysis we have identified patterns and groups that are of particular interest for bacterial and archaeal phages. Even though only a small subset of phages was classified at the order and family level, we observed interesting trends that are worth exploring. At the order level, the orders Vinavirales, Kalamavirales, Durnavirales, Asfuvirales, Patatavirales, Zurhausenvirales and Ortervirales were only found in bacterial phages. In the same fashion, the familes *Corticoviridae, Peduoviridae, Casjensviridae, Tectiviridae, Roundtreeviridae, Duneviridae, Partitiviridae, Zierdtviridae, Chaseviridae, Hytrosaviridae, Guelinviridae, Asfarviridae, Steigviridae, Salasmaviridae, Papilloviridae, Potyviridae* and *Caullimoviridae* were only found in bacterial phages. However, we observed some unexpected families: *Lipothrixviridae* and *Rudiviridae* in bacteria; these families have only been reported in archaeal hosts^46^.

At the family level, archaeal phages also demonstrated exclusivity as *Globuloviridae* and *Malacoherpesviridae* were only found in the archaea group. The presence of *Globuloviridae* is expected, given that only archaeal hosts have been reported^47^. Conversely, the detection of *Malacoherpesviridae* is unexpected, as it is known to infect bivalves^48^. Interestingly, the family *Retroviridae*, which is found in vertebrate hosts^49^, was found infecting unclassified hosts. These identified phages that are exclusive to particular groups could have unique signatures that might shed some light in differences between archaeal and bacterial phages. However, it is possible that these phages simply haven’t been discovered across these groups yet but could be found with more sampling.

Additional findings of interest include the environmental distribution of the prophage hosts. It is noteworthy that the marine source was the most prevalent in archaeal and unclassified samples, while it was the least prevalent in bacterial samples. This might suggest that human-related environments are not major reservoirs for prophages, or it could simply indicate a sampling bias which is the most likely. For example, in activated sludge, phages are highly abundant and diverse and more viral DNA has been found in this environment in comparison with soil and plant-associated environments, and other engineered systems^50^.

Ultimately, these results emphasize the need for sampling in new environments and regions, which could lead to the discovery of novel microbial species and their prophages. Additionally, there is a need to improve metadata collection practices, as our database lacks environmental information for some entries. Prophage-DB can enable finer analyses of prophage distributions since it has been demonstrated that some phages are globally distributed while others are endemic^51^. Therefore, it would be worth investigating if there are genetic underpinnings for what makes a phage endemic or not.

In our prophage count analysis of Prophage-DB, we identified host groups of interest which contained high numbers of prophages. At the phylum level, in bacteria, Pseudomonadota was the predominant group in both prophage and host counts and has been reported in other studies^52^. Other groups with high prophage densities include Firmicutes and Actinobacteria, likely from a high degree of sampling due to their relevance in medical research^53^.

For each taxonomic rank, we identified over and under-represented groups. Generally, our database had more phage diversity compared to IMG/VR v4, and thus several groups were not included in this analysis as they were not found in IMG/VR v4 (**Table S7-S12**). The unclassified category tended to be over-represented across all ranks suggesting a higher portion of novel phages in our database in comparison to those reported in IMG/VR v4 (this was observed across anthropogenic, freshwater, host-associated, marine, and terrestrial environments). In bacterial hosts, phage taxonomic groups for the most part were over-represented or were not over or under-represented. In contrast, archaeal phages tended to be underrepresented across all taxonomic ranks. Studying these viruses is essential for enhancing viral sequence databases, as this research will likely contribute to the discovery of additional viruses and the refinement of alignment-independent methods.

In our analysis of auxiliary metabolic genes, we examined how they are distributed across several environments and hosts. It has been hypothesized that AMG compositions may reflect the adaptation of their hosts to their environments^24^, that AMGs show a niche-dependent distribution pattern^31^ and that some environments show a higher AMG diversity^54^. Our comparison of AMGs across the selected environments did not reveal any noticeable differences which presents a contrast to previous reports. Regarding AMG distribution across hosts, we looked in the top 20 most abundant phyla and did not observe any stark contrast across them. Similarly to AMG environmental distribution, host distribution was mostly uniform but there were certain phyla with specific groups of AMGs being more abundant. This suggests that AMG composition differences might be more influenced by the host.

## CONCLUSION

Prophage-DB is an extensive database encompassing over 350,000 prophage sequences from archaeal and bacterial hosts. Prophage-DB provides a multifaceted exploration of prophages, examining their relationships with their hosts, their environmental distribution, their auxiliary metabolic genes (AMGs), and their taxonomy. Prophage-DB is highly representative in comparison to virus databases. Given the fundamental role of prophages in microbiomes, Prophage-DB will enable studies of prophage diversity, ecology, and evolution in microbiomes. We anticipate that Prophage-DB will serve as a valuable resource for virus and phage research, offering insights into prophages and their roles in microbiomes and ecosystems.

## Data availability

The prophage genomic sequences (.fna files) are available in the DRYAD repository alongside metadata: https://doi.org/10.5061/dryad.3n5tb2rs5

## Code availability

Codes used in this project are available at the following GitHub repository: https://github.com/AnantharamanLab/Prophage-DB

### Acknowledgments

This research was supported by National Institute of General Medical Sciences of the National Institutes of Health under award number R35GM143024. ED was supported by the NIH Biotechnology Traineeship under Award Number T32GM135066. CM was supported by an NSF Graduate Research Fellowship and a UW-Madison SciMed GRS Fellowship.

## Author contributions

ED and KA conceived the project. ED conducted bioinformatic analyses, statistical analyses, visualization of results, and content organization. CM conducted AMG and statistical analyses. ED and KA wrote the manuscript draft. All authors reviewed the results, edited, and approved the manuscript.

## Competing interests

The authors declare no competing interests.

## Supplementary information

<u>Additional file 1: Supplementary Table 1 (A, B, C)</u>

File format: .xlsb

Title of data: Metadata associated with prophage genomes Description of data: The first sheet contains an explanation of each column. Data has been split into three files.

<u>Additional file 2: Supplementary Table 2-24</u>

File format: .xlsb

Title of data: AMG metadata, Comparison of GTDB (release 220) taxonomy to observed taxonomic ranks with prophages in Prophage-DB, CheckV summary results, Prophage counts, Representation analysis counts, Comparison of observed and expected counts

Description of data: The first sheet contains an explanation of each sheet listed.

<u>Additional file 3: Supplementary Figures 1-24</u>

File format: .docx

Title of data: Counts for taxonomic ranks, prophage counts, representation analysis plots, Comparison of observed and expected counts

Description of data: This file contains supplementary figures 1-24.

## Notes

https://doi.org/10.5061/dryad.3n5tb2rs5

### Competing Interest Statement

The authors have declared no competing interest.

