## Supplementary Figures 1-24 for "Prophage-DB: A comprehensive database to explore diversity, distribution, and ecology of prophages"

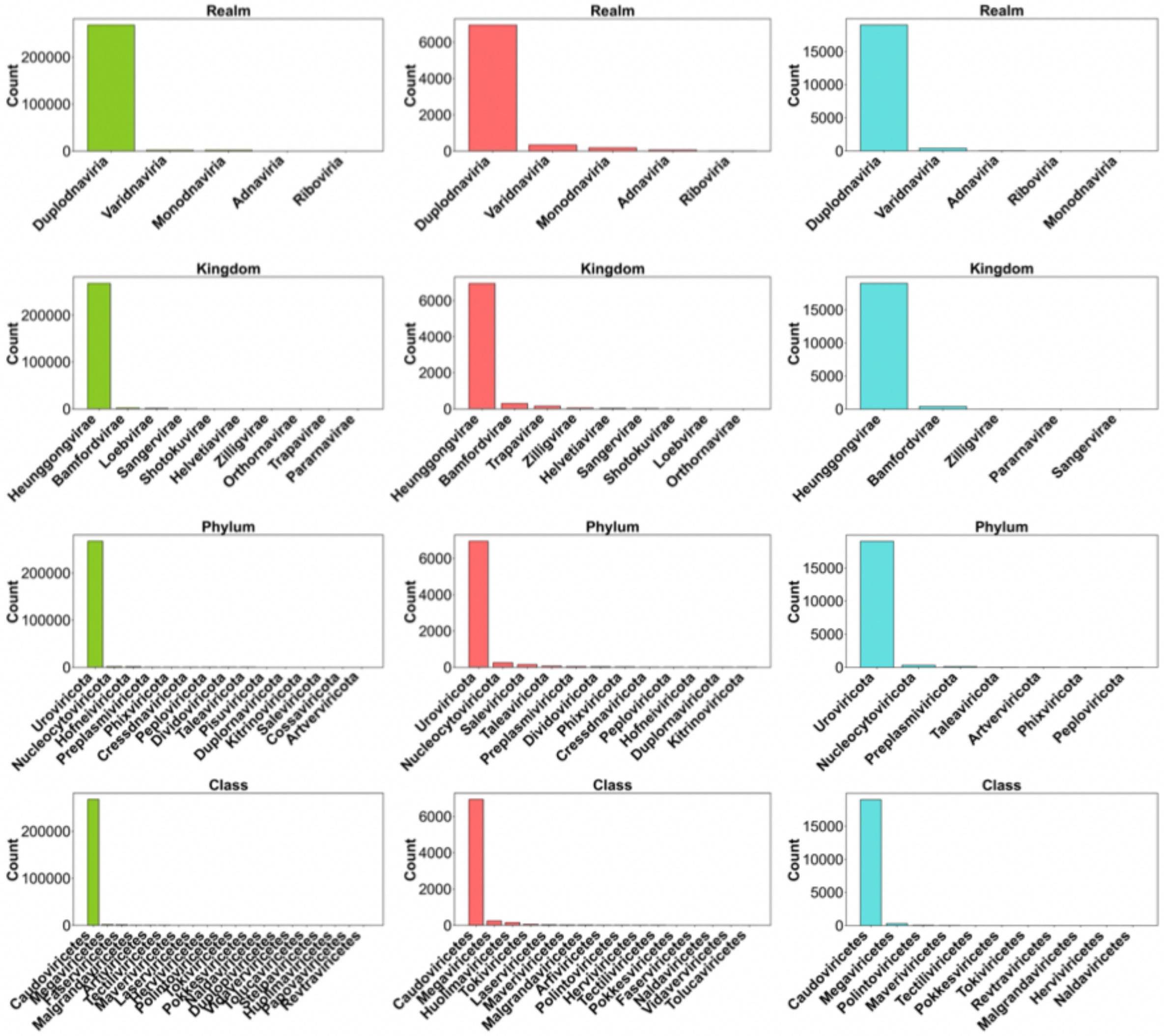


**Supplementary figure 1** Counts for taxonomic ranks (realm, kingdom, phylum, and class). Bacterial phages (green), archaeal phages (red), phages from unclassified hosts (blue).

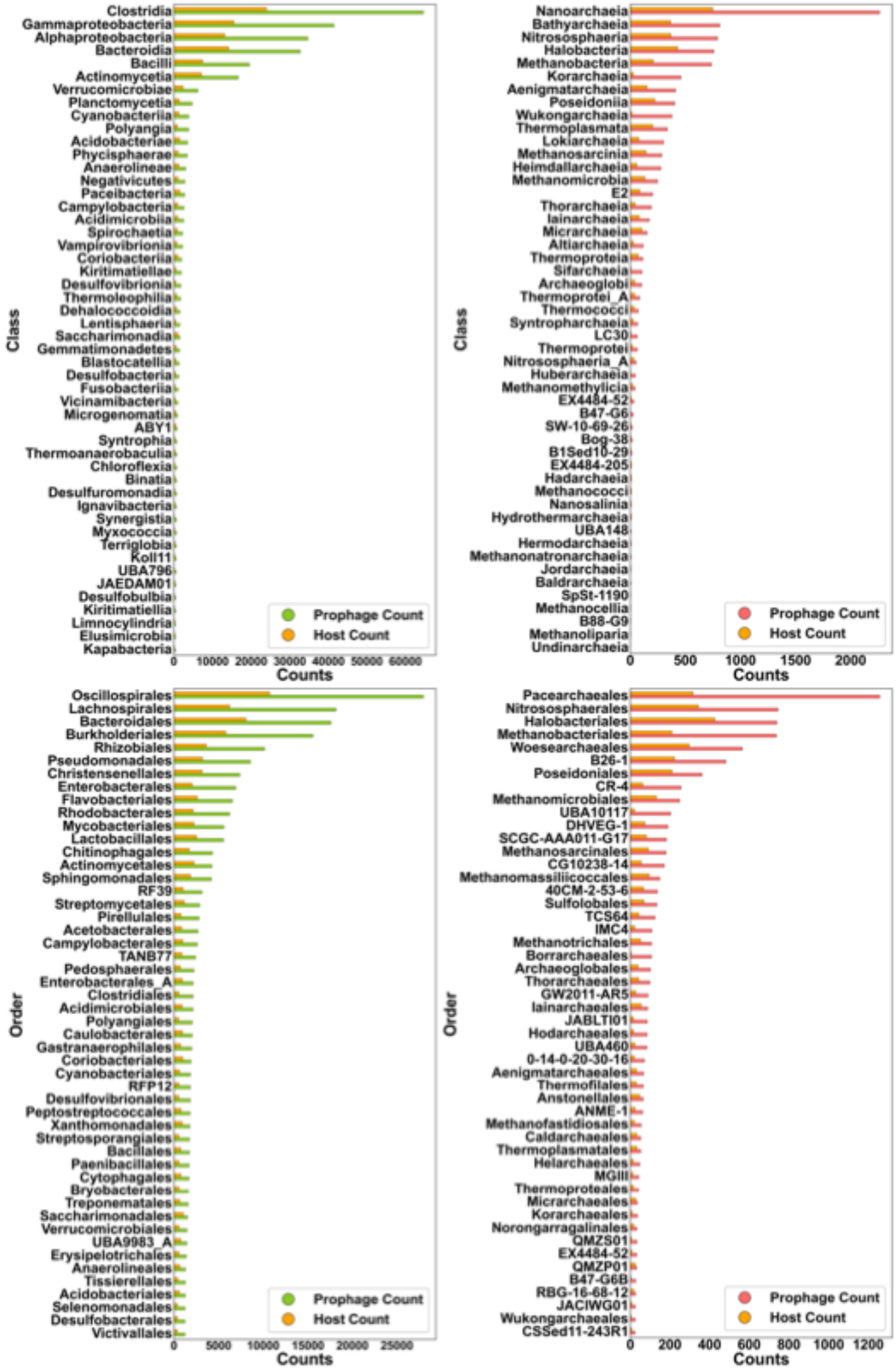


**Supplementary figure 2** Plots with host and prophage counts for bacteria (left side plots) and archaea (right side plots). Plots are only showing the first 50 taxonomic groups with more prophage counts at the class and order levels.

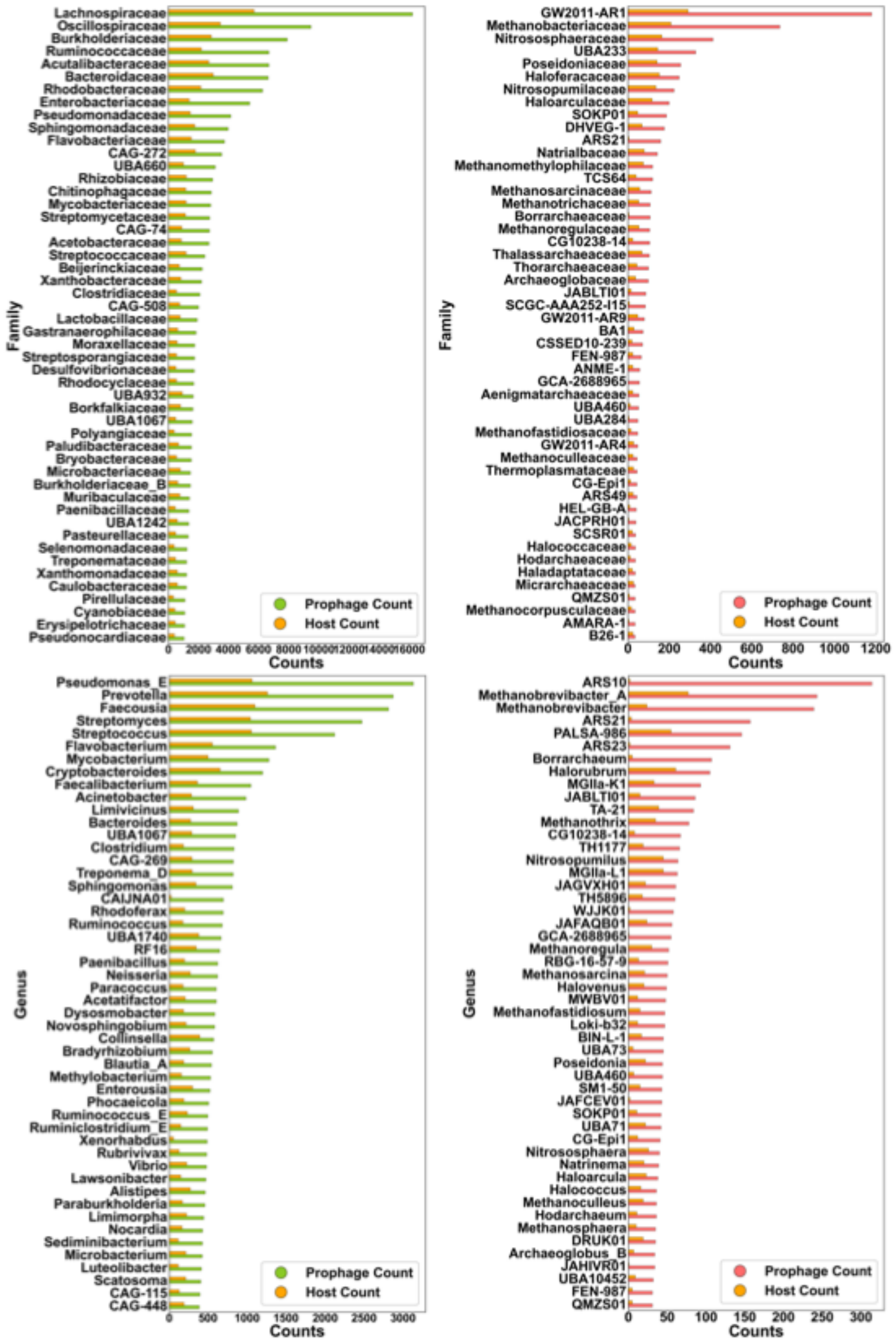


**Supplementary figure 3** Plots with host and prophage counts for bacteria (left side plots) and archaea (right side plots). Plots are only showing the first 50 taxonomic groups with more prophage counts at the family and genus levels.

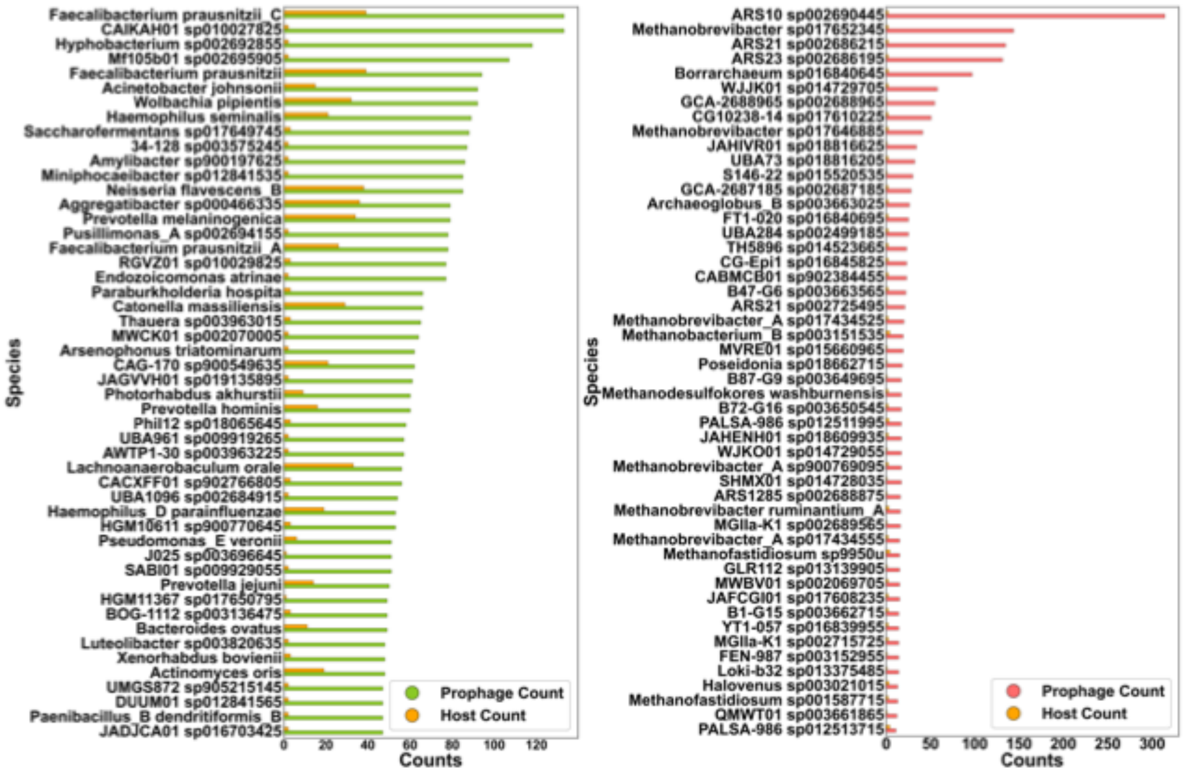


**Supplementary figure 4** Plots with host and prophage counts for bacteria (left side plots) and archaea (right side plots). Plots are only showing the first 50 taxonomic groups with more prophage counts at the species level.

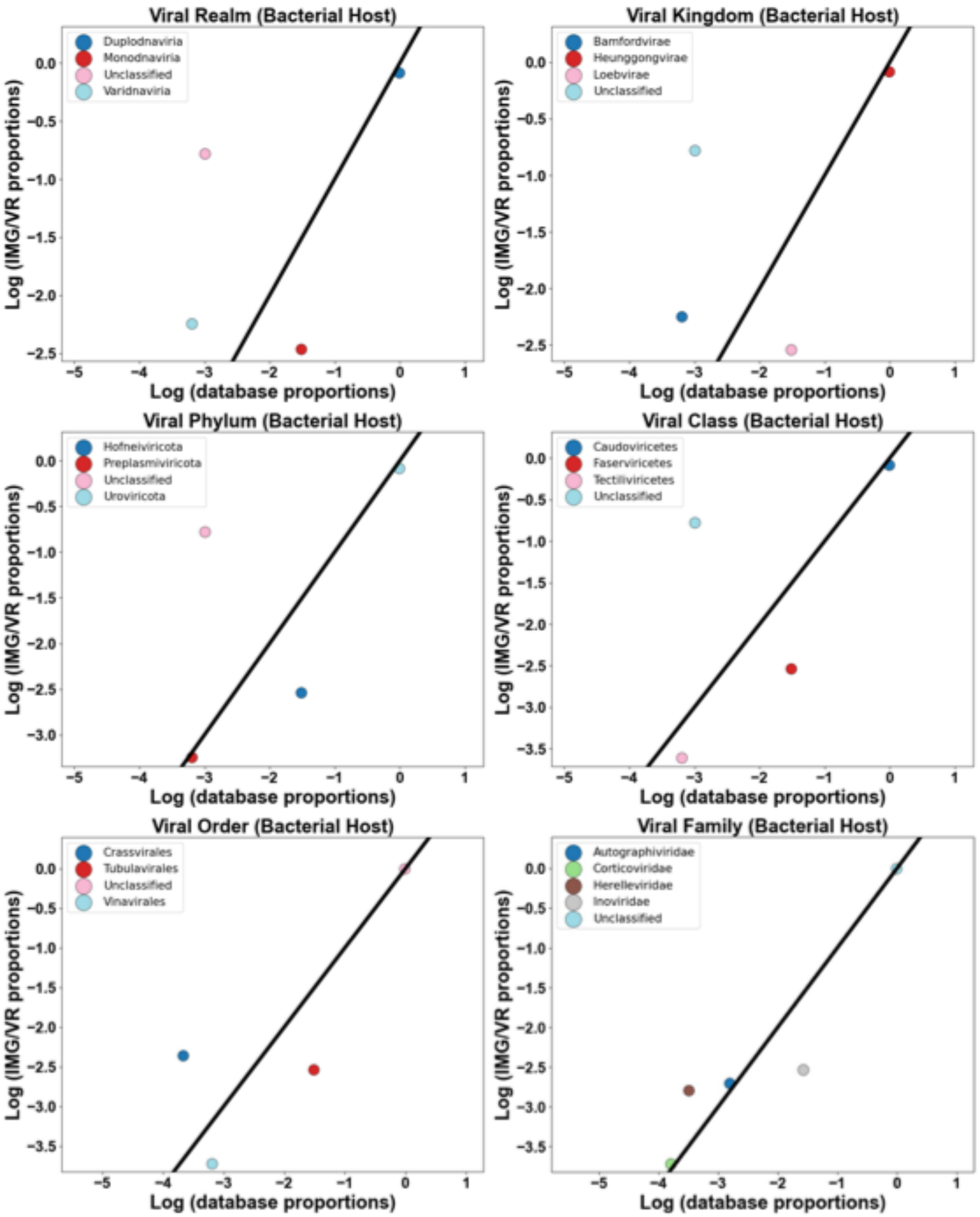


**Supplementary figure 5.** Representation analysis comparing proportion of taxonomic groups in our database to the proportions in IMG/VR database. Using log transformed values. Values above the 1:1 line are considered over-represented, those below, are under-represented. Data shows phage representation in bacterial hosts in anthropogenic environments.

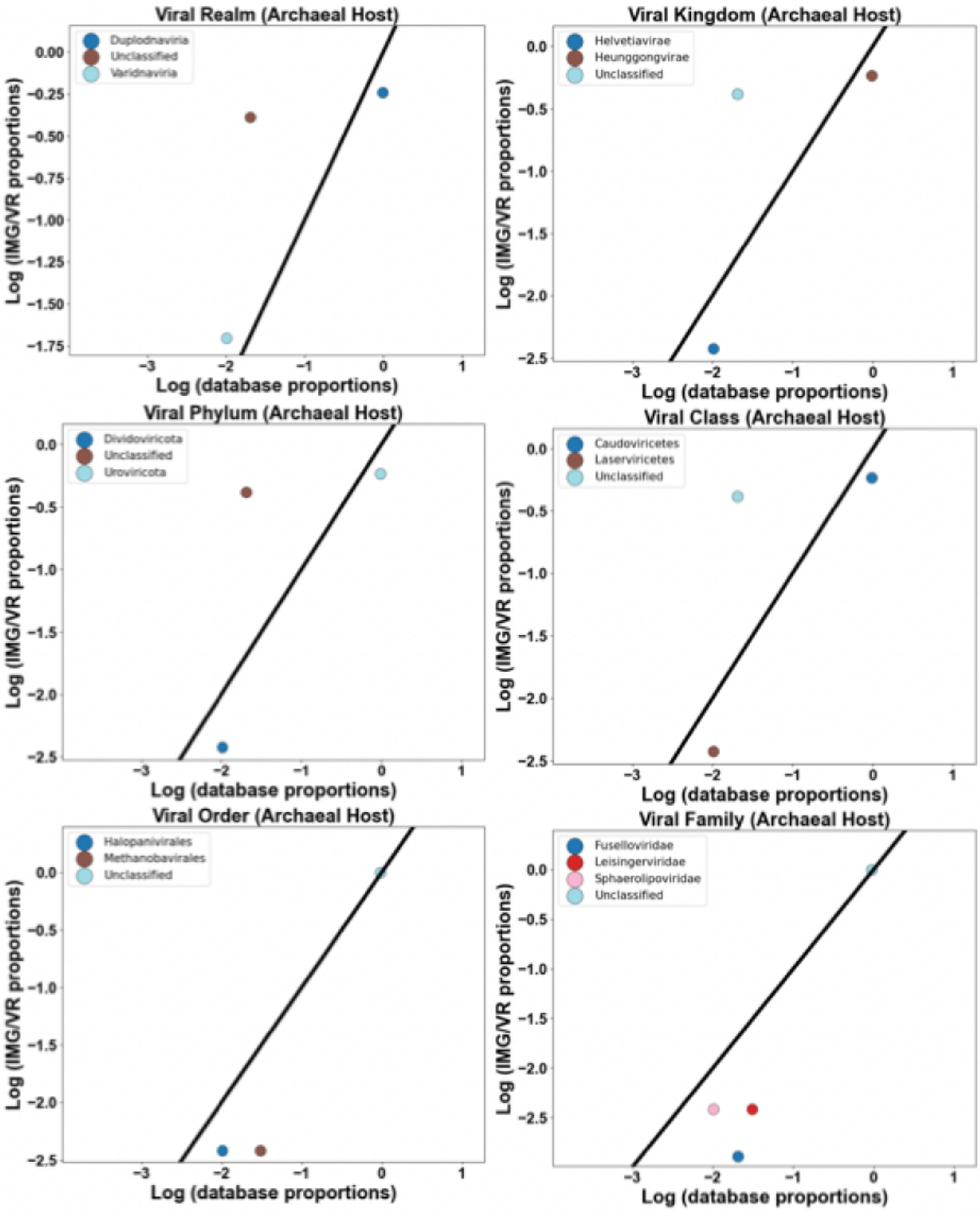


**Supplementary figure 6.** Representation analysis comparing proportion of taxonomic groups in our database to the proportions in IMG/VR database. Using log transformed values. Values above the 1:1 line are considered over-represented, those below, are under-represented. Data shows phage representation in archaeal hosts in anthropogenic environments.

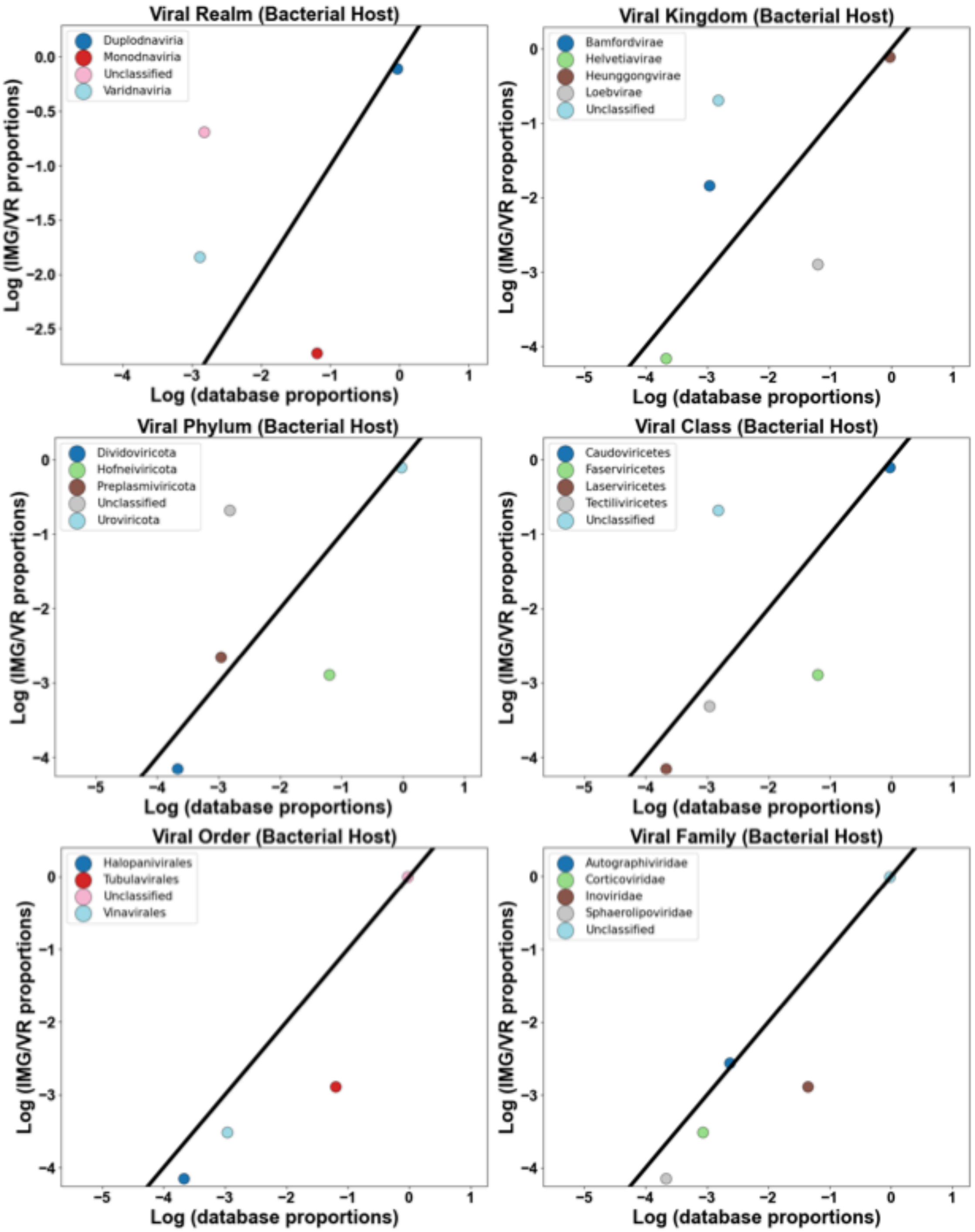


**Supplementary figure 7.** Representation analysis comparing proportion of taxonomic groups in our database to the proportions in IMG/VR database. Using log transformed values. Values above the 1:1 line are considered over-represented, those below, are under-represented. Data shows phage representation in bacterial hosts in freshwater environments.

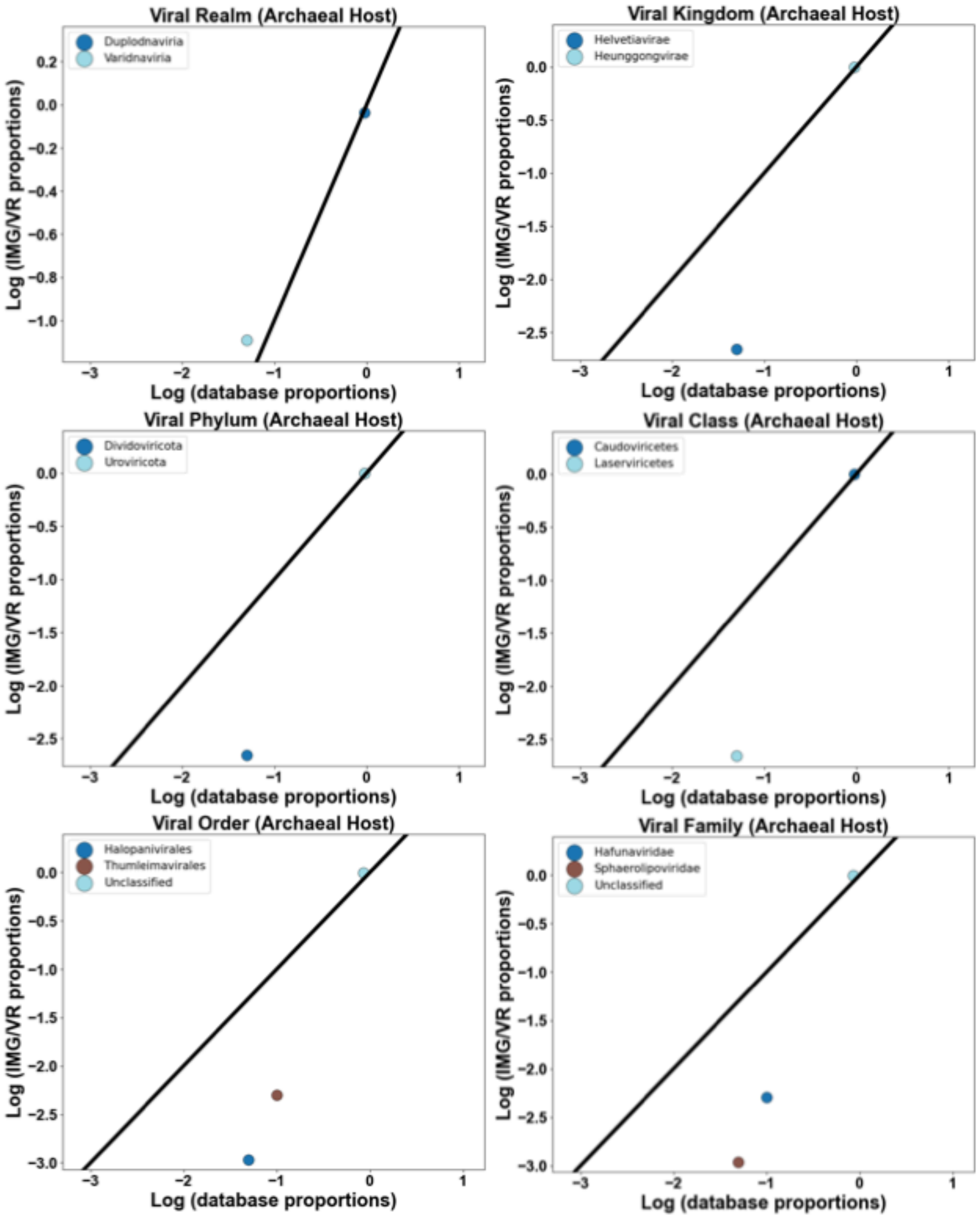


**Supplementary figure 8.** Representation analysis comparing proportion of taxonomic groups in our database to the proportions in IMG/VR database. Using log transformed values. Values above the 1:1 line are considered over-represented, those below, are under-represented. Data shows phage representation in archaeal hosts in freshwater environments.

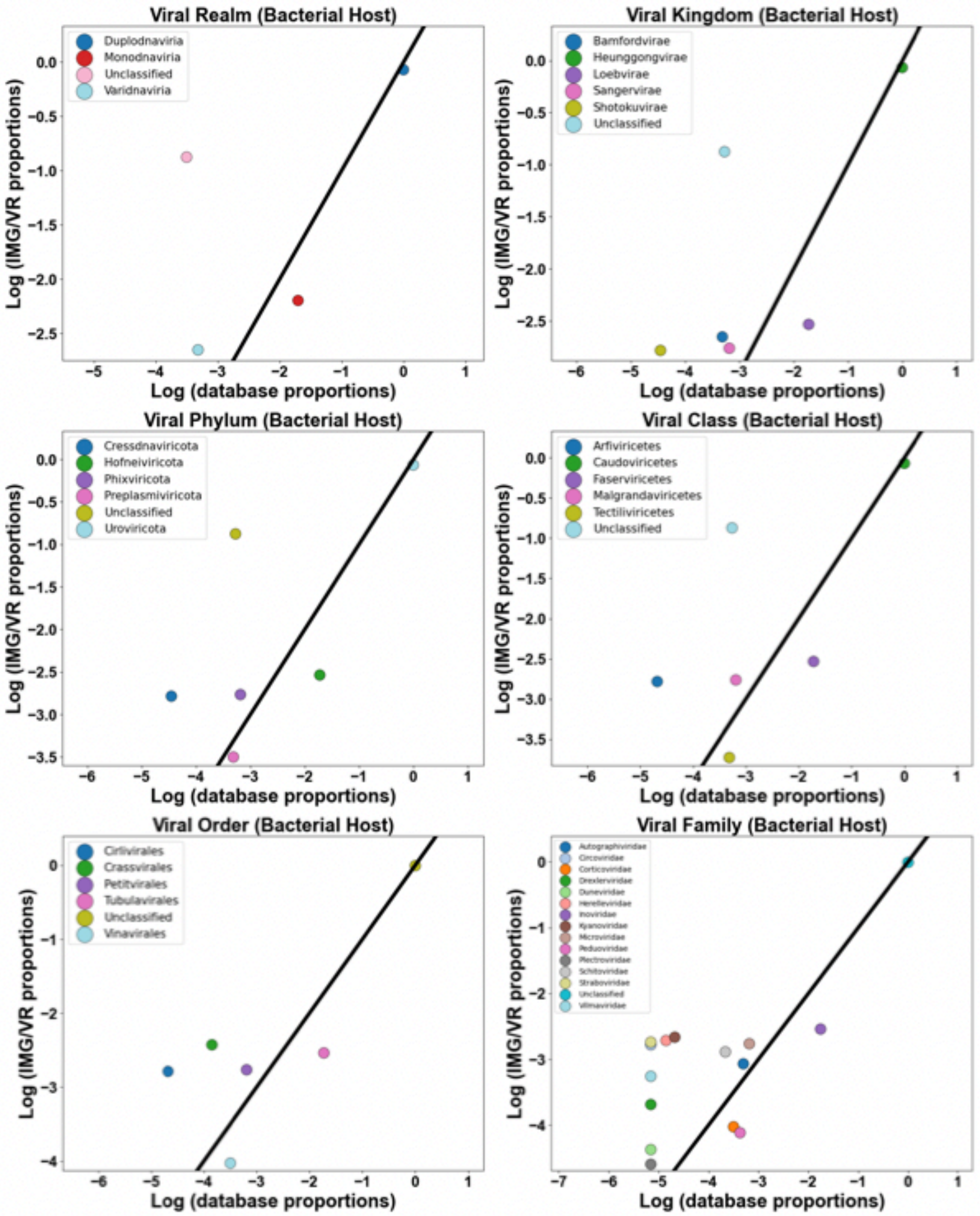


**Supplementary figure 9.** Representation analysis comparing proportion of taxonomic groups in our database to the proportions in IMG/VR database. Using log transformed values. Values above the 1:1 line are considered over-represented, those below, are under-represented. Data shows phage representation in bacterial hosts in host-associated environments.

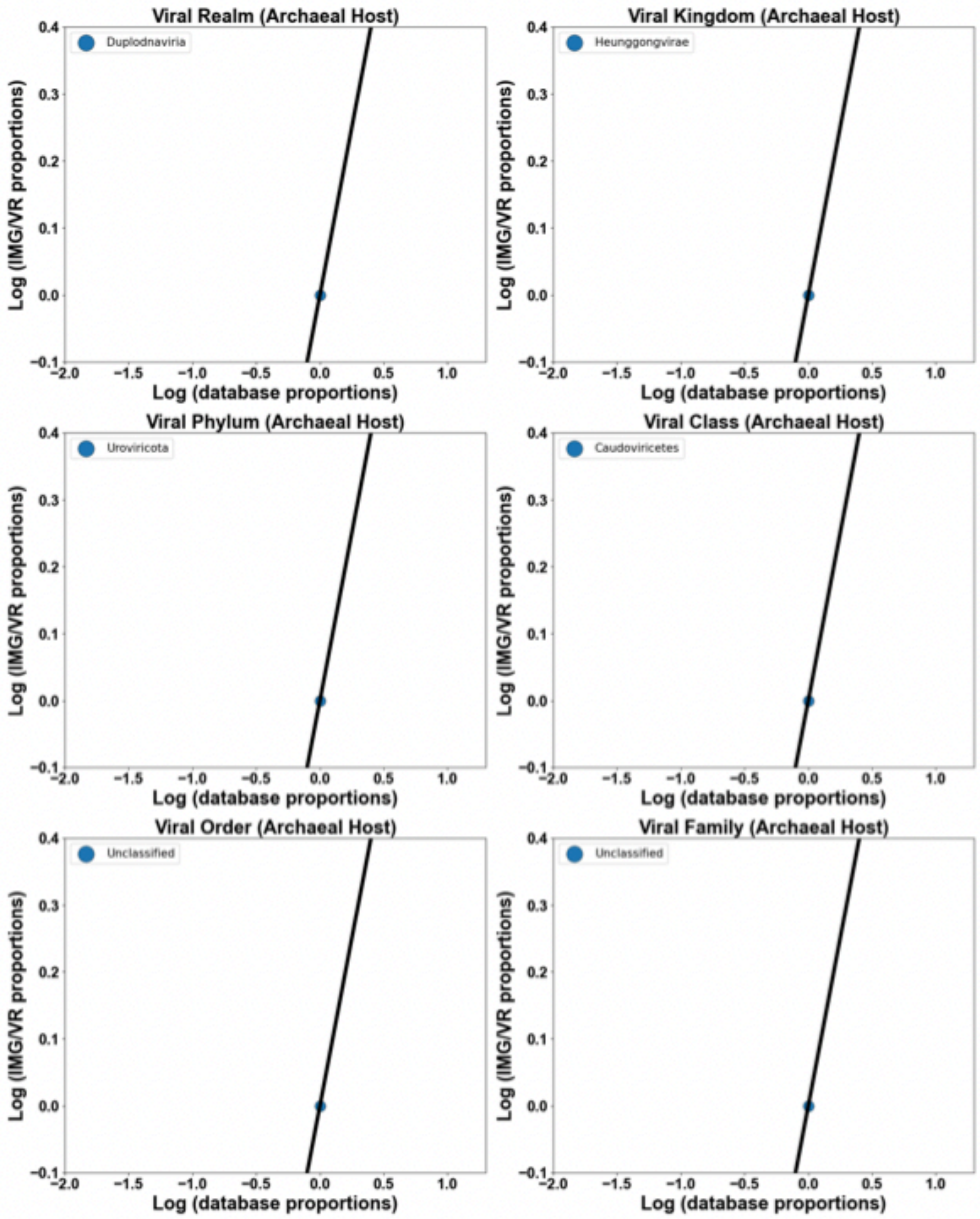


**Supplementary figure 10.** Representation analysis comparing proportion of taxonomic groups in our database to the proportions in IMG/VR database. Using log transformed values. Values above the 1:1 line are considered over-represented, those below, are under-represented. Data shows phage representation in archaeal hosts in host-associated environments.

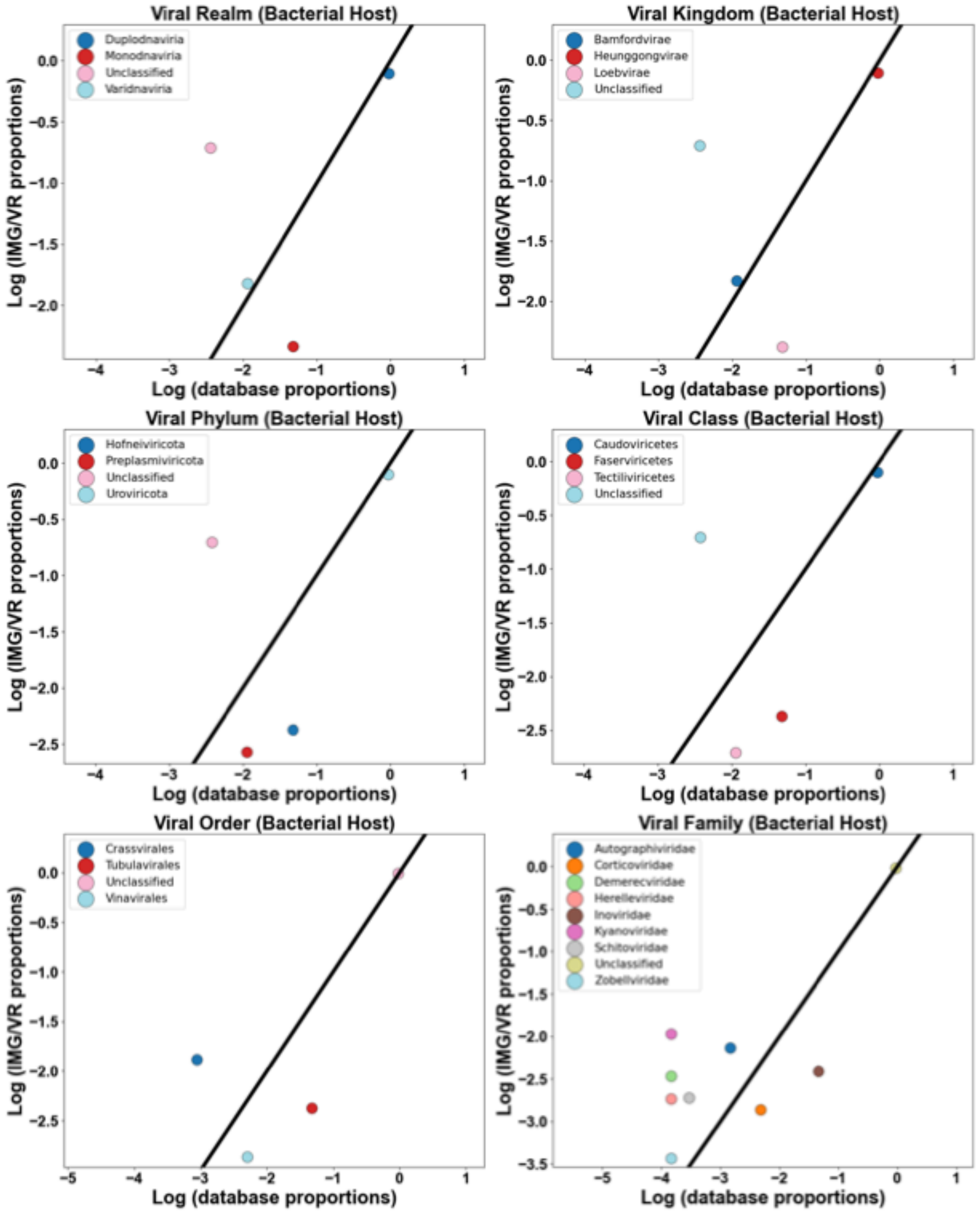


**Supplementary figure 11.** Representation analysis comparing proportion of taxonomic groups in our database to the proportions in IMG/VR database. Using log transformed values. Values above the 1:1 line are considered over-represented, those below, are under-represented. Data shows phage representation in bacterial hosts in marine environments.

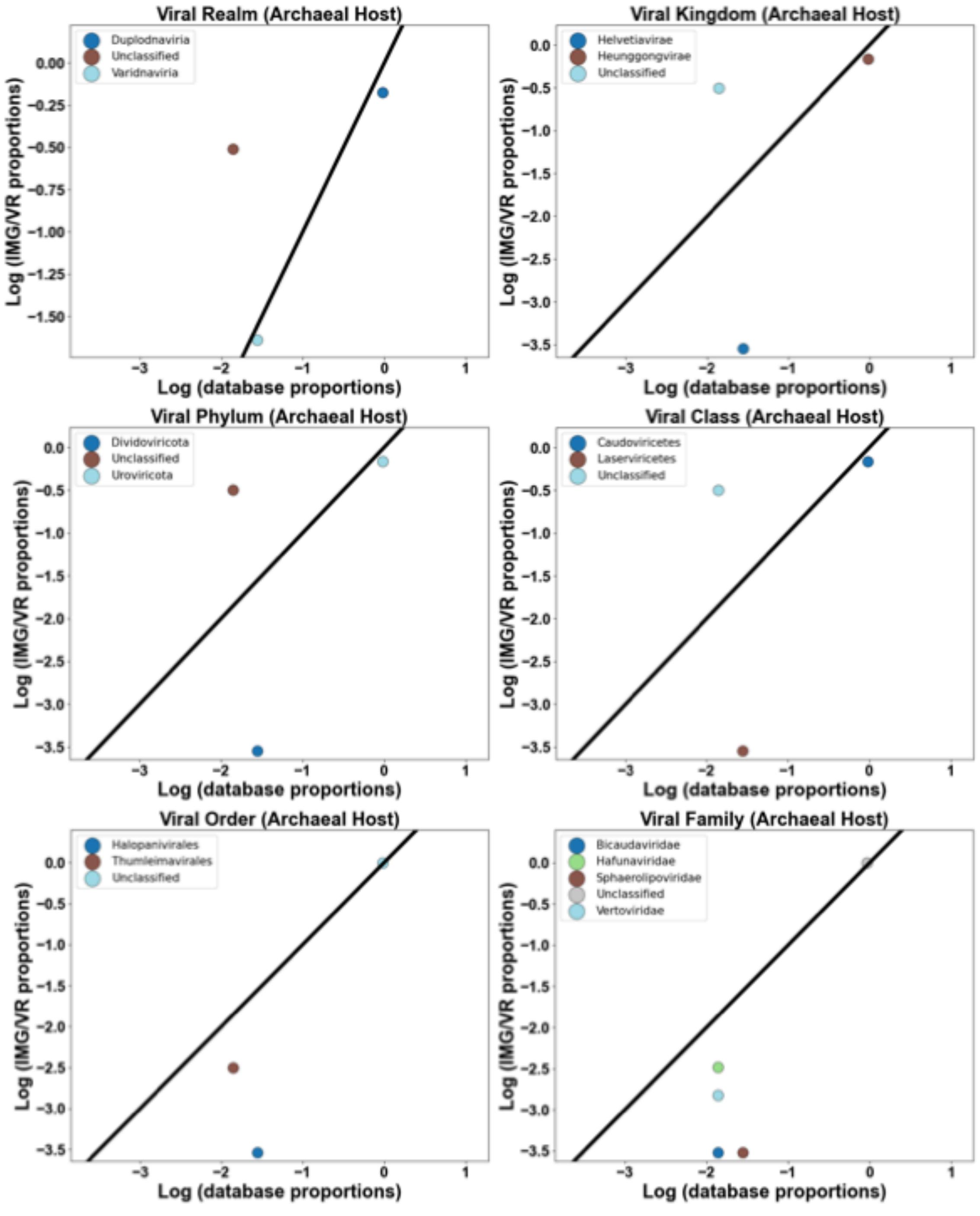


**Supplementary figure 12.** Representation analysis comparing proportion of taxonomic groups in our database to the proportions in IMG/VR database. Using log transformed values. Values above the 1:1 line are considered over-represented, those below, are under-represented. Data shows phage representation in archaeal hosts in marine environments.

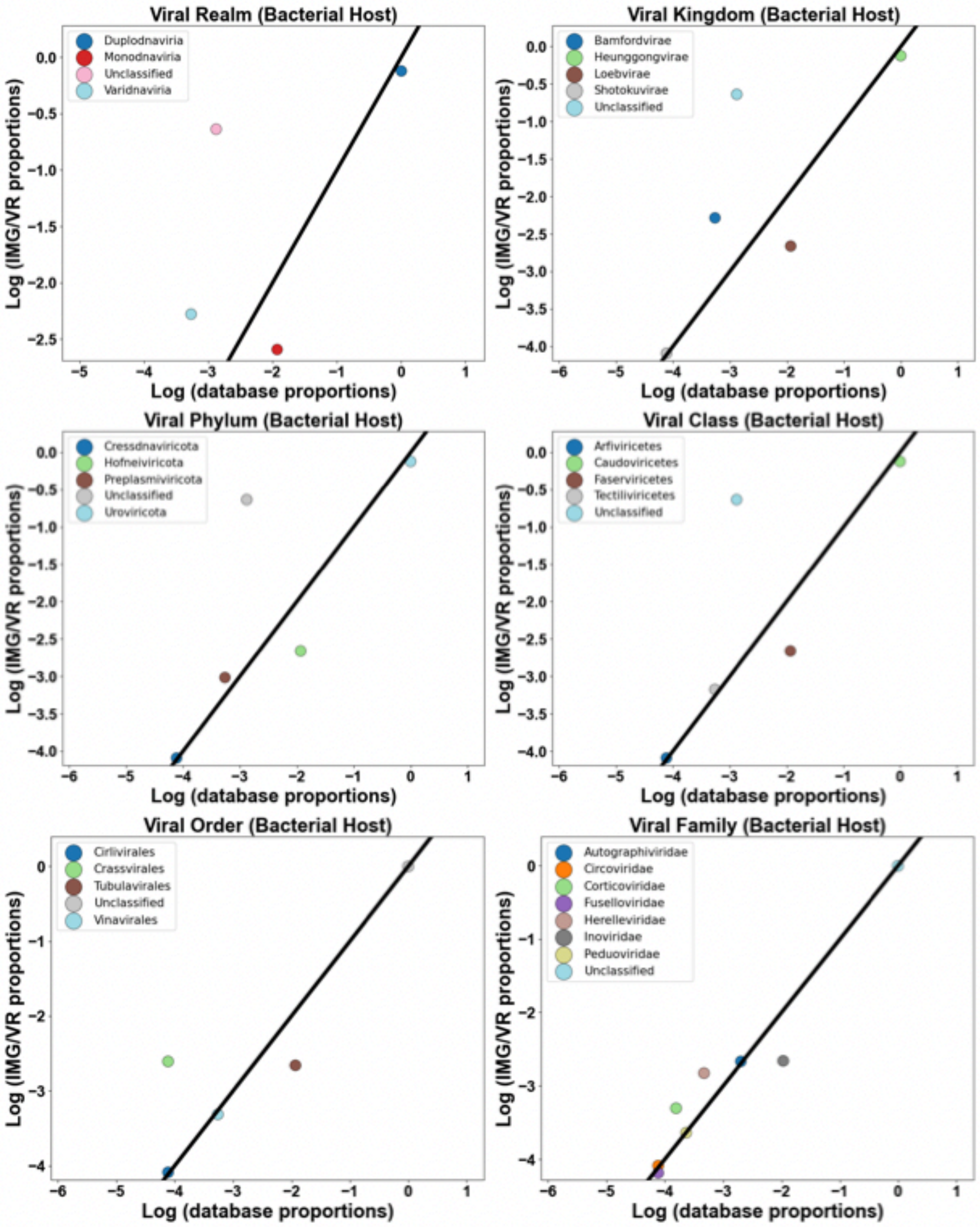


**Supplementary figure 13.** Representation analysis comparing proportion of taxonomic groups in our database to the proportions in IMG/VR database. Using log transformed values. Values above the 1:1 line are considered over-represented, those below, are under-represented. Data shows phage representation in bacterial hosts in terrestrial environments.

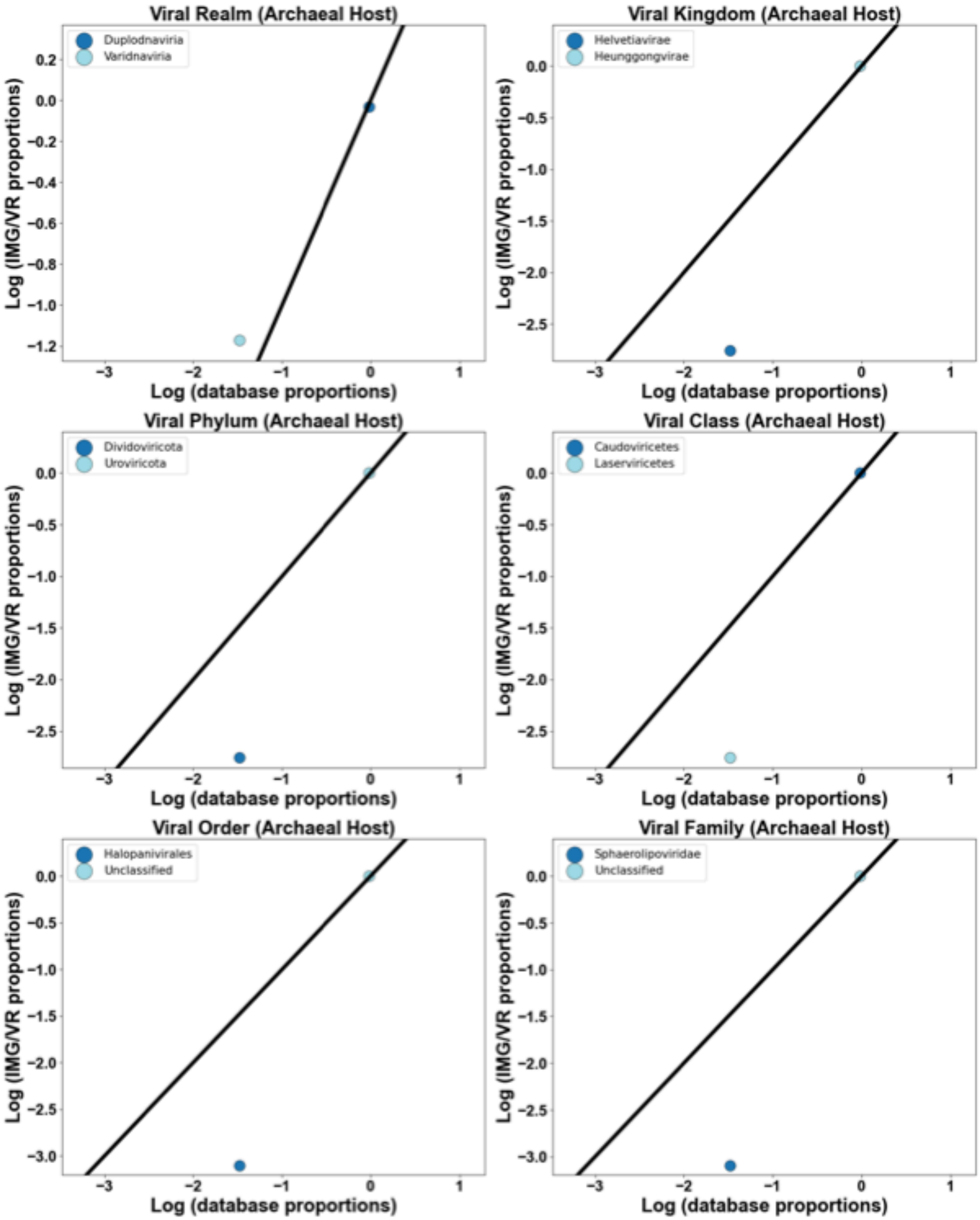


**Supplementary figure 14.** Representation analysis comparing proportion of taxonomic groups in our database to the proportions in IMG/VR database. Using log transformed values. Values above the 1:1 line are considered over-represented, those below, are under-represented. Data shows phage representation in archaeal hosts in terrestrial environments.

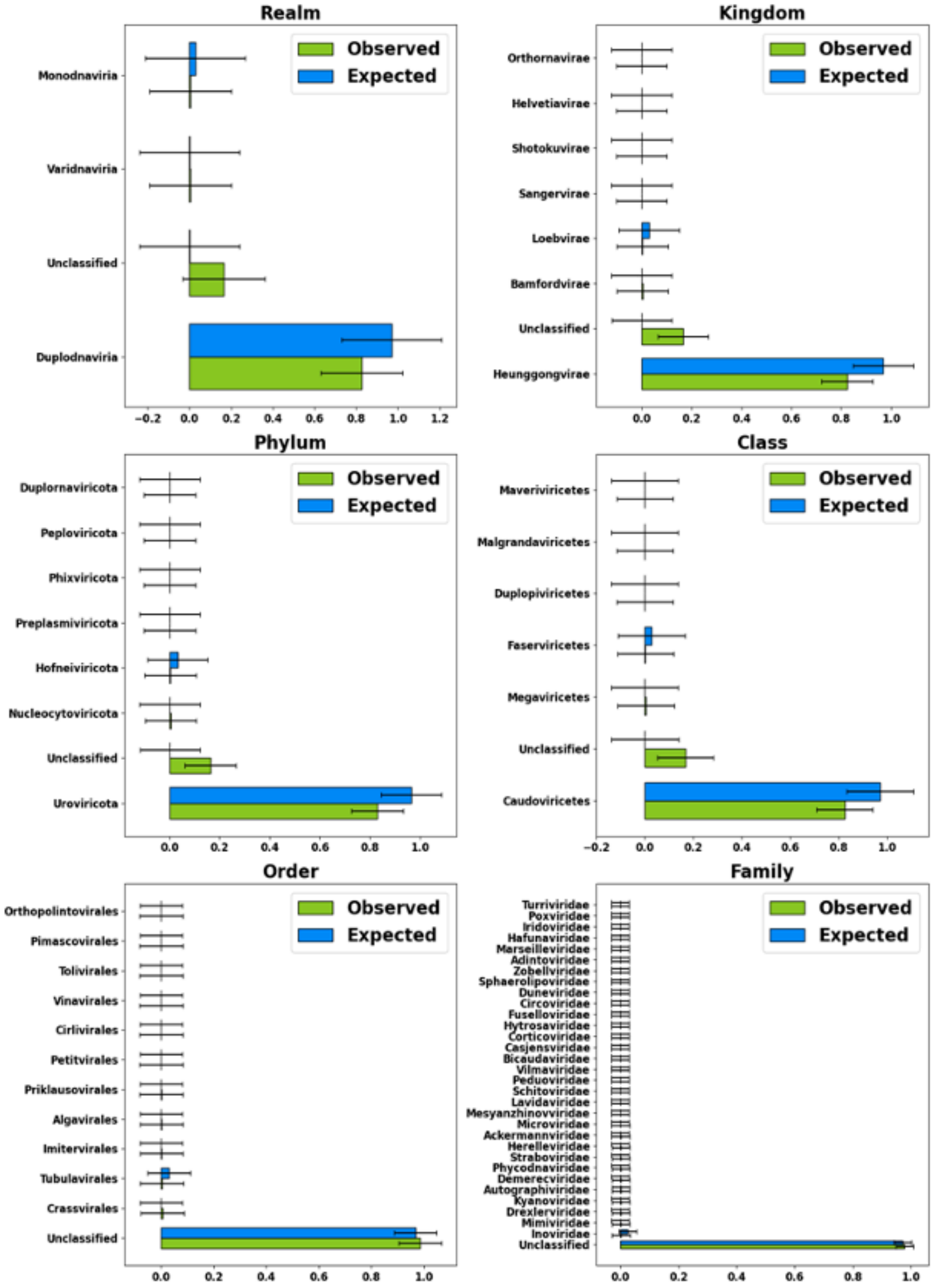


**Supplementary figure 15.** Comparison between IMG/VR counts (expected) and Prophage-DB counts (observed) for bacterial phages in anthropogenic environments. Random sampling with replacement was used (bootstrap value of 10,000).

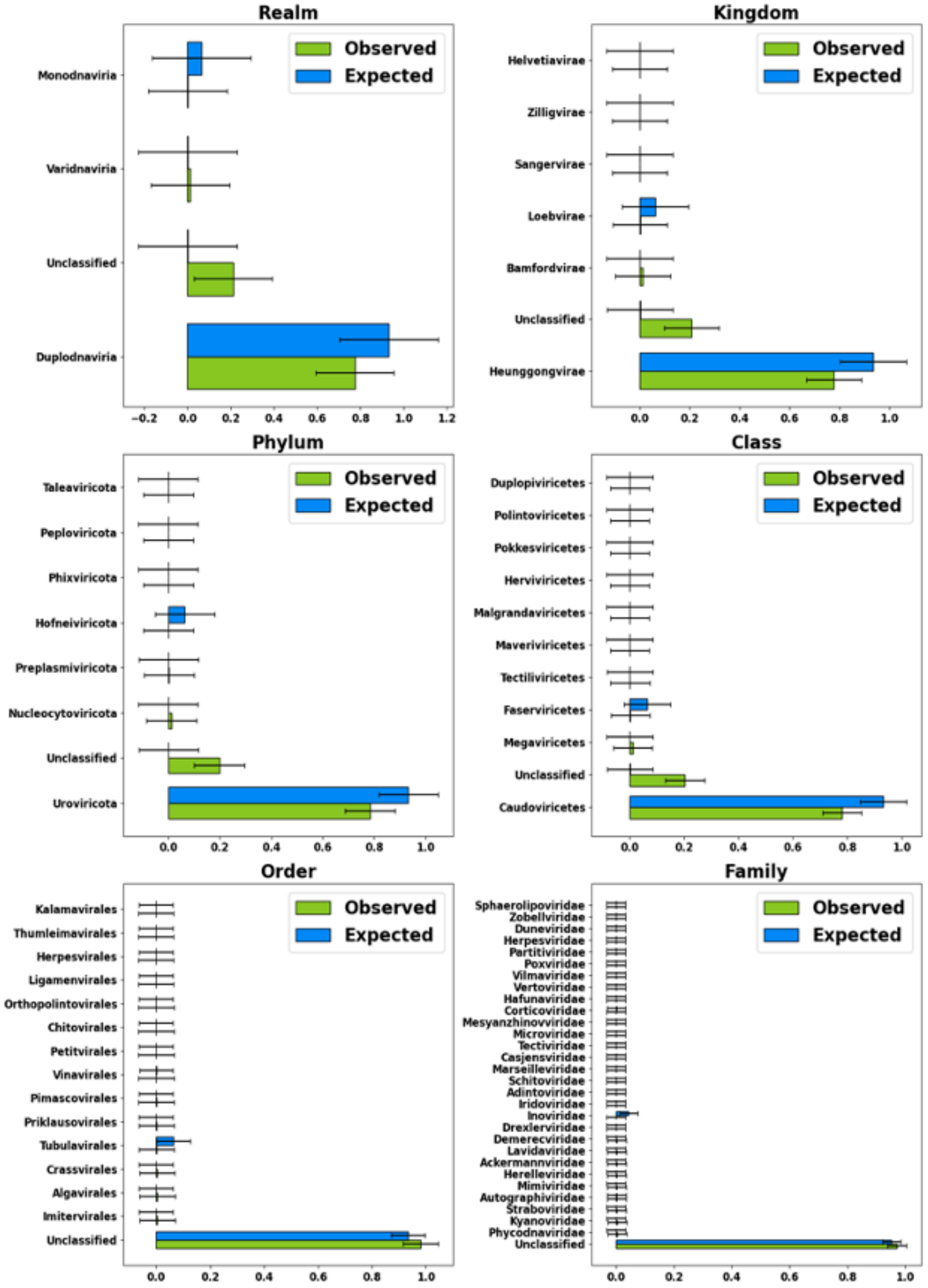


**Supplementary figure 16.** Comparison between IMG/VR counts (expected) and Prophage-DB counts (observed) for bacterial phages in freshwater environments. Random sampling with replacement was used (bootstrap value of 10,000).

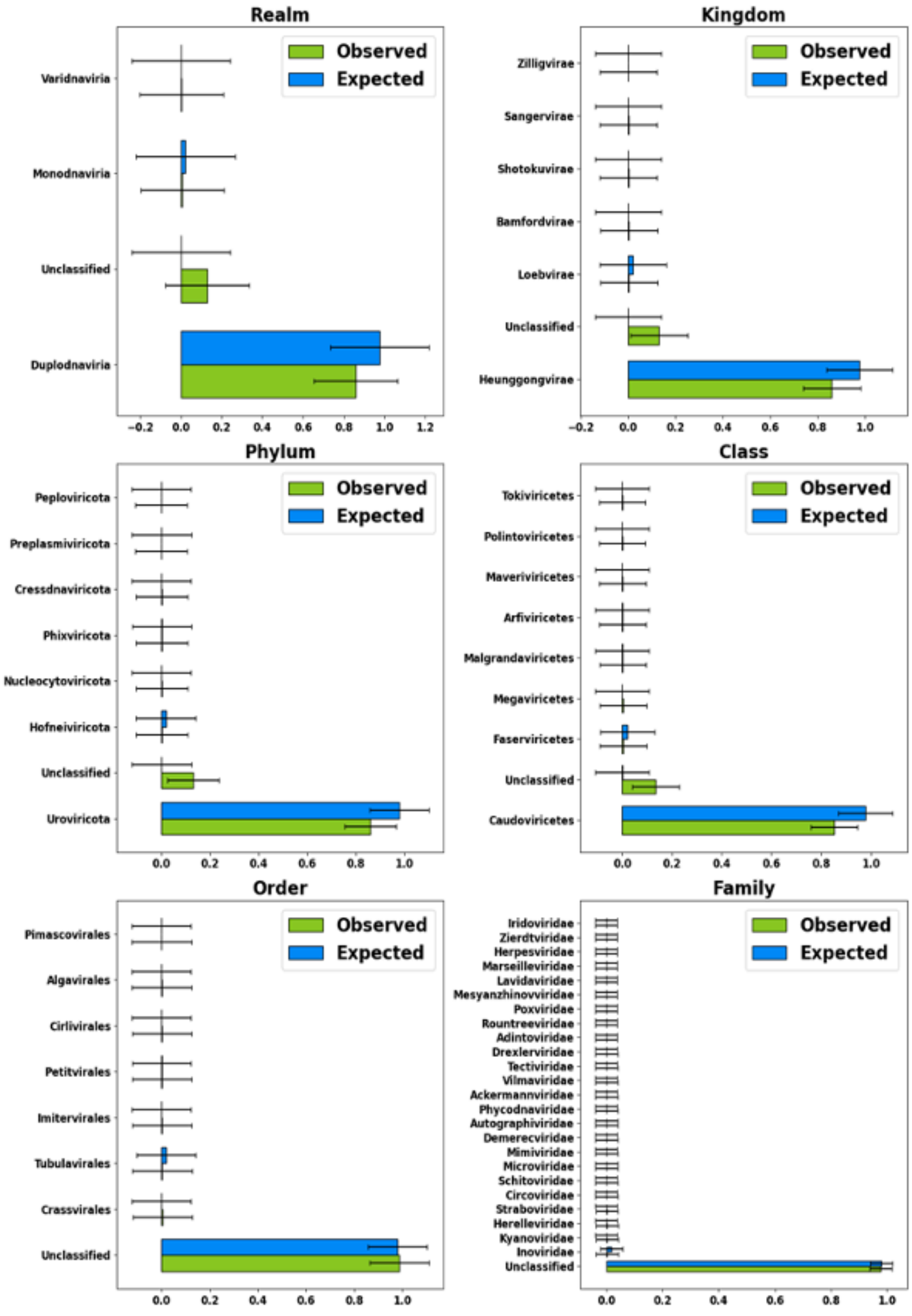


**Supplementary figure 17.** Comparison between IMG/VR counts (expected) and Prophage-DB counts (observed) for bacterial phages in host-associated environments. Random sampling with replacement was used (bootstrap value of 10,000).

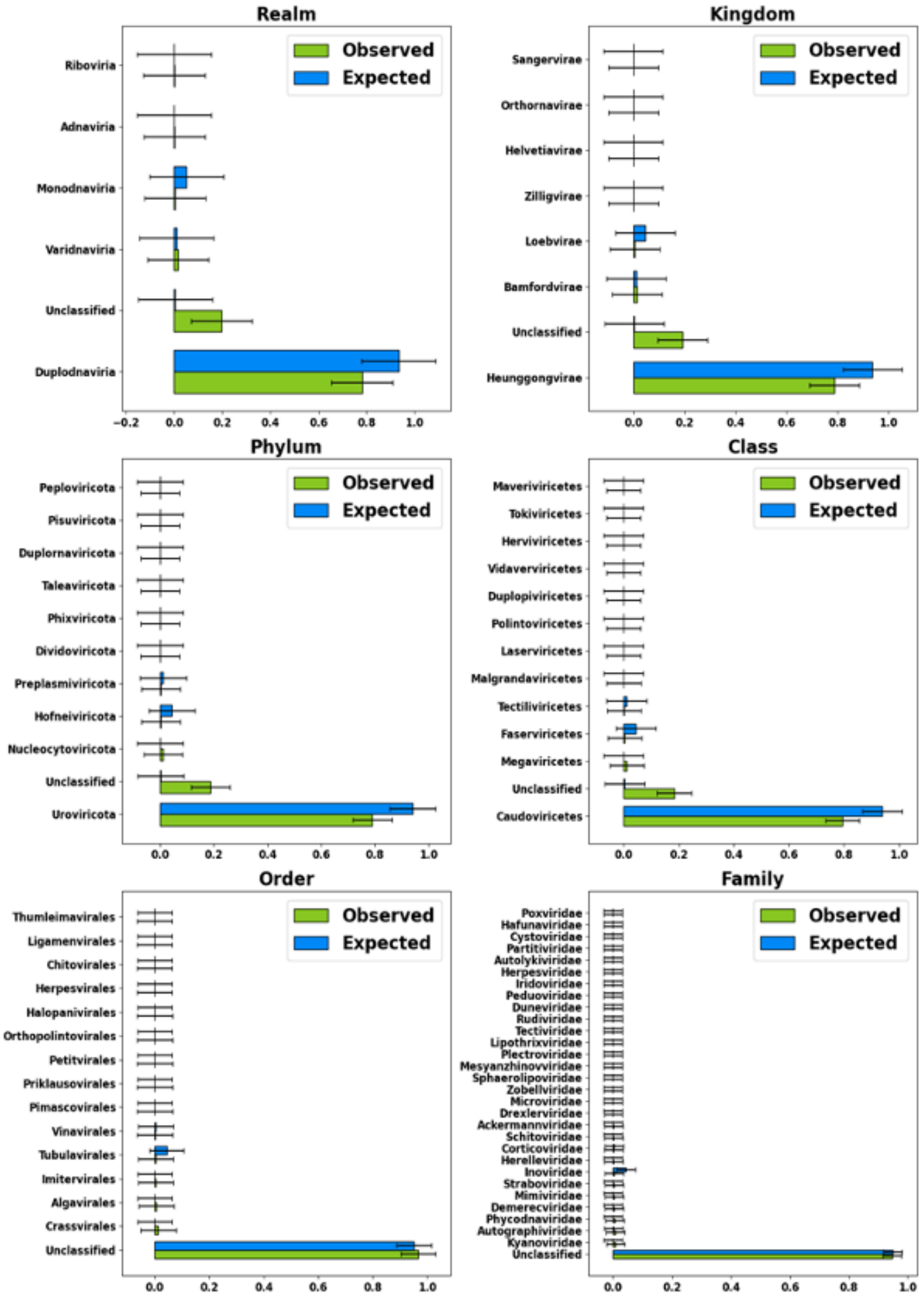


**Supplementary figure 18.** Comparison between IMG/VR counts (expected) and Prophage-DB counts (observed) for bacterial phages in marine environments. Random sampling with replacement was used (bootstrap value of 10,000).

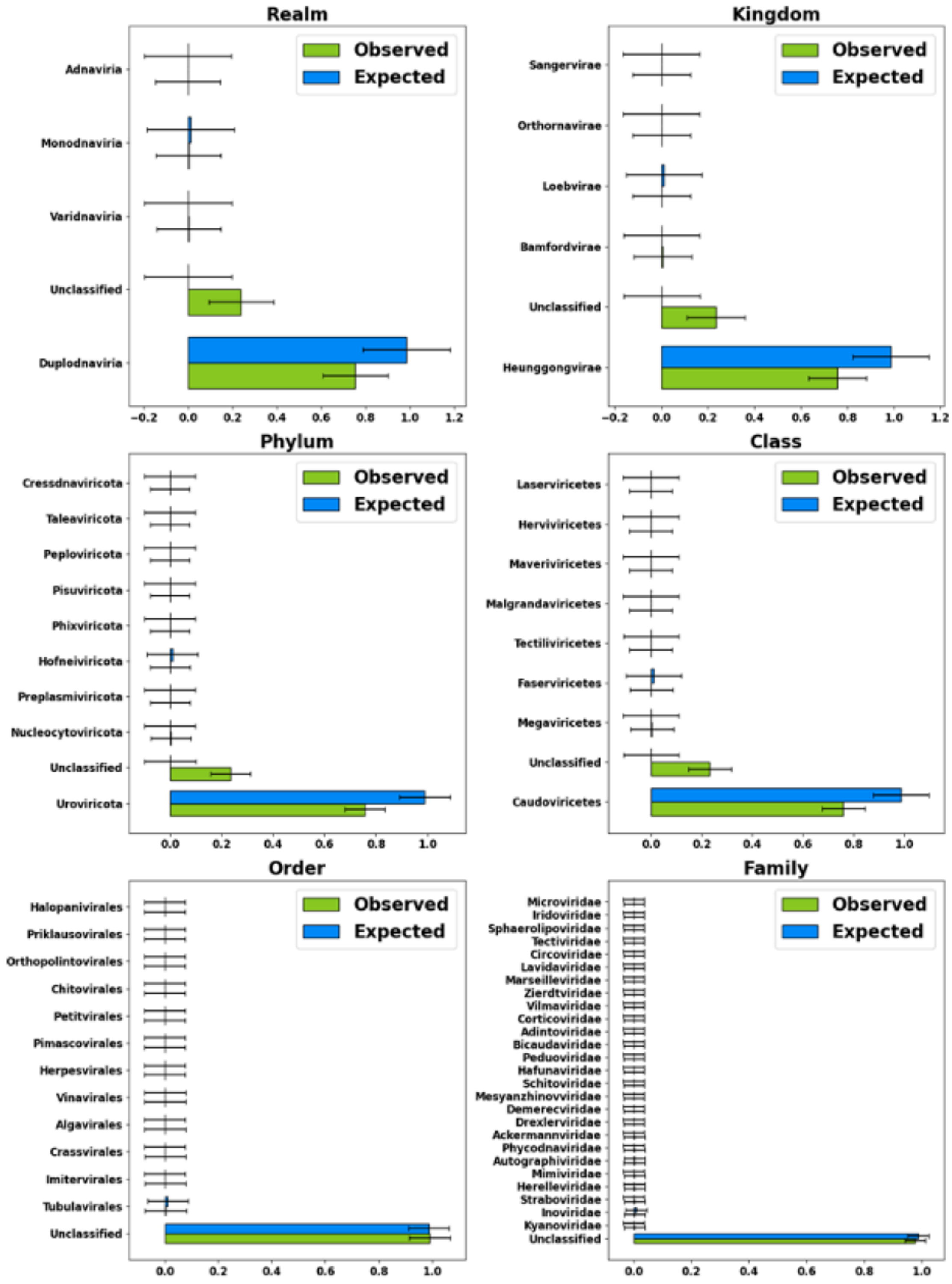


**Supplementary figure 19.** Comparison between IMG/VR counts (expected) and Prophage-DB counts (observed) for bacterial phages in terrestrial environments. Random sampling with replacement was used (bootstrap value of 10,000).

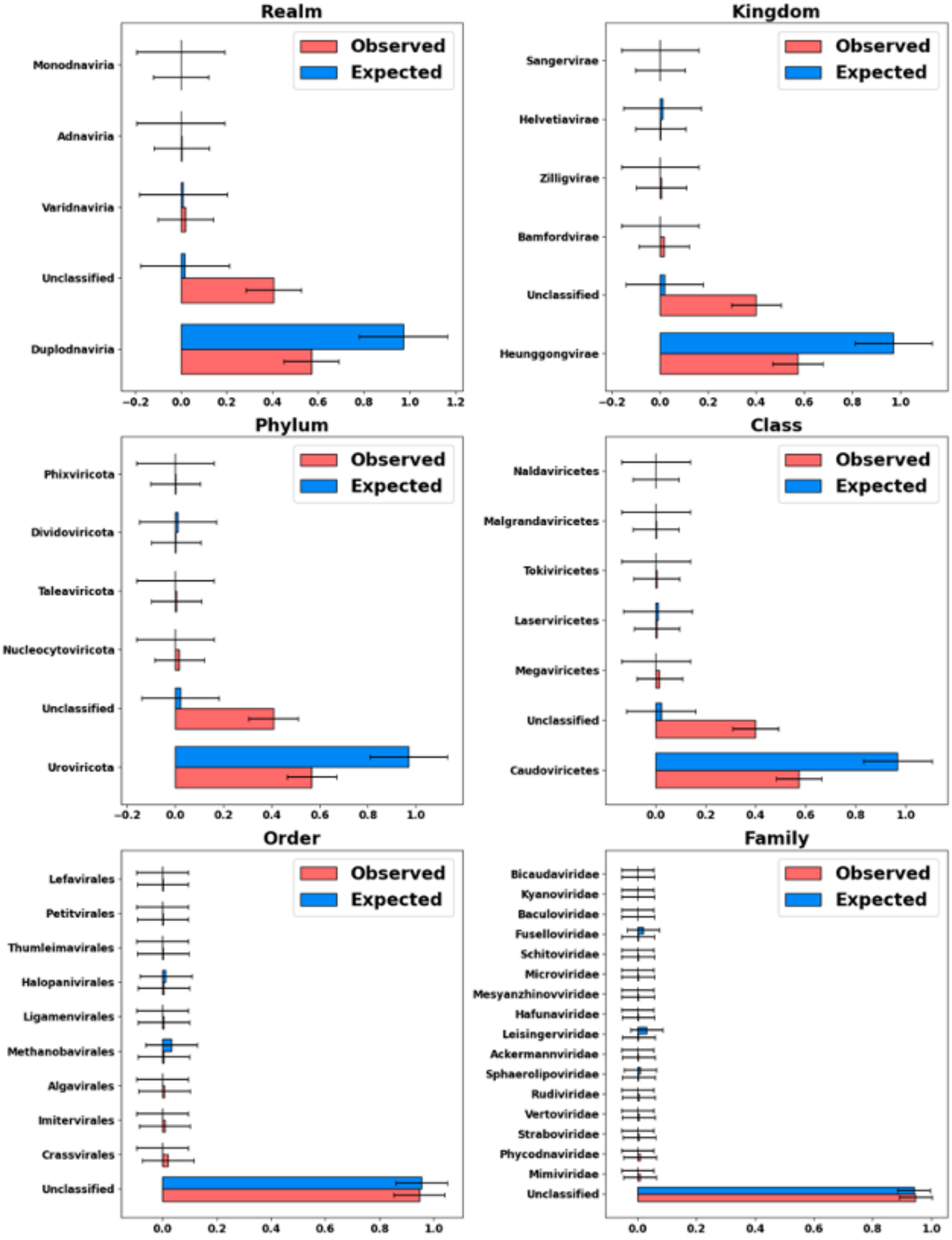


**Supplementary figure 20.** Comparison between IMG/VR counts (expected) and Prophage-DB counts (observed) for archaeal phages in anthropogenic environments. Random sampling with replacement was used (bootstrap value of 10,000).

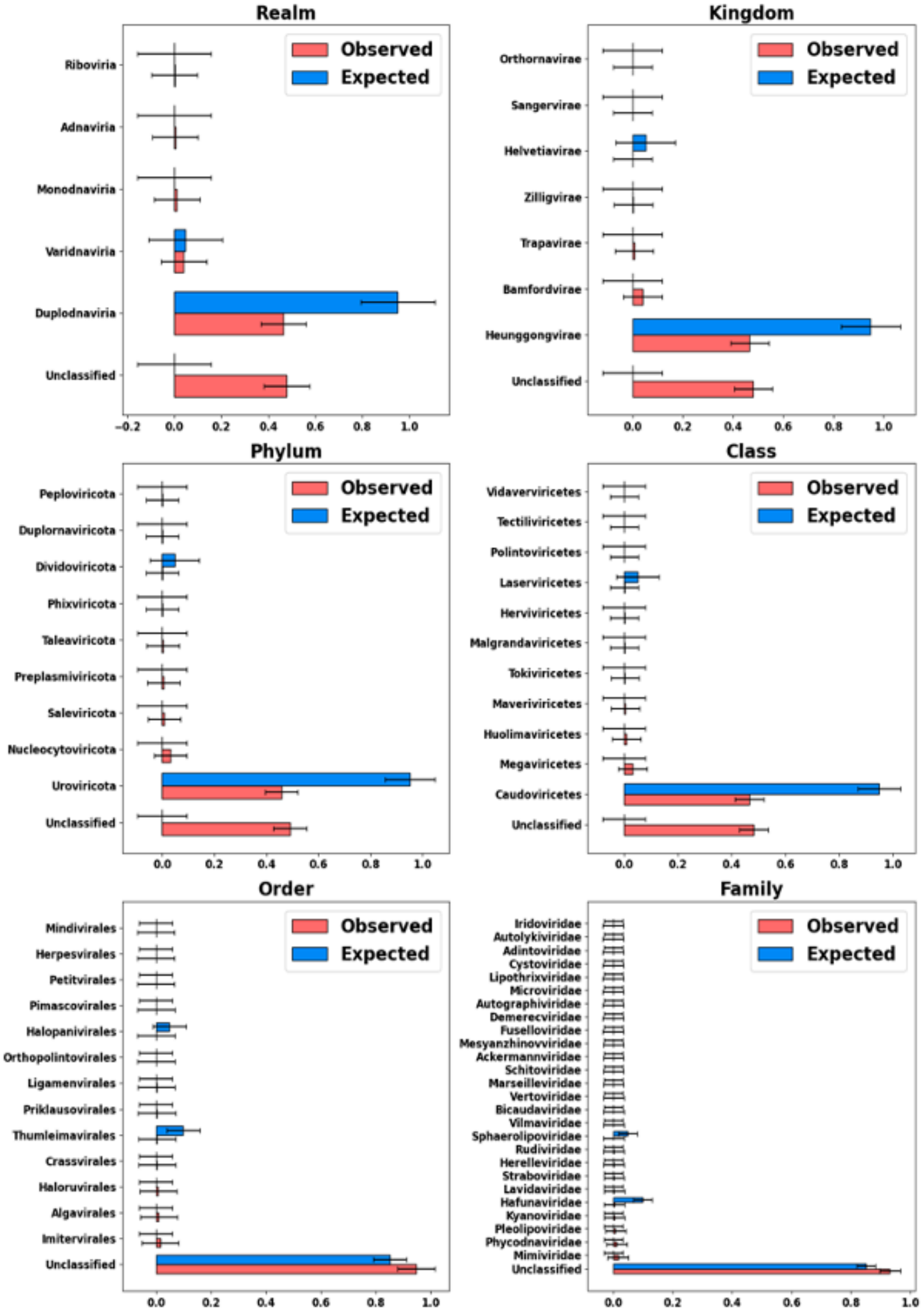


**Supplementary figure 21.** Comparison between IMG/VR counts (expected) and Prophage-DB counts (observed) for archaeal phages in freshwater environments. Random sampling with replacement was used (bootstrap value of 10,000).

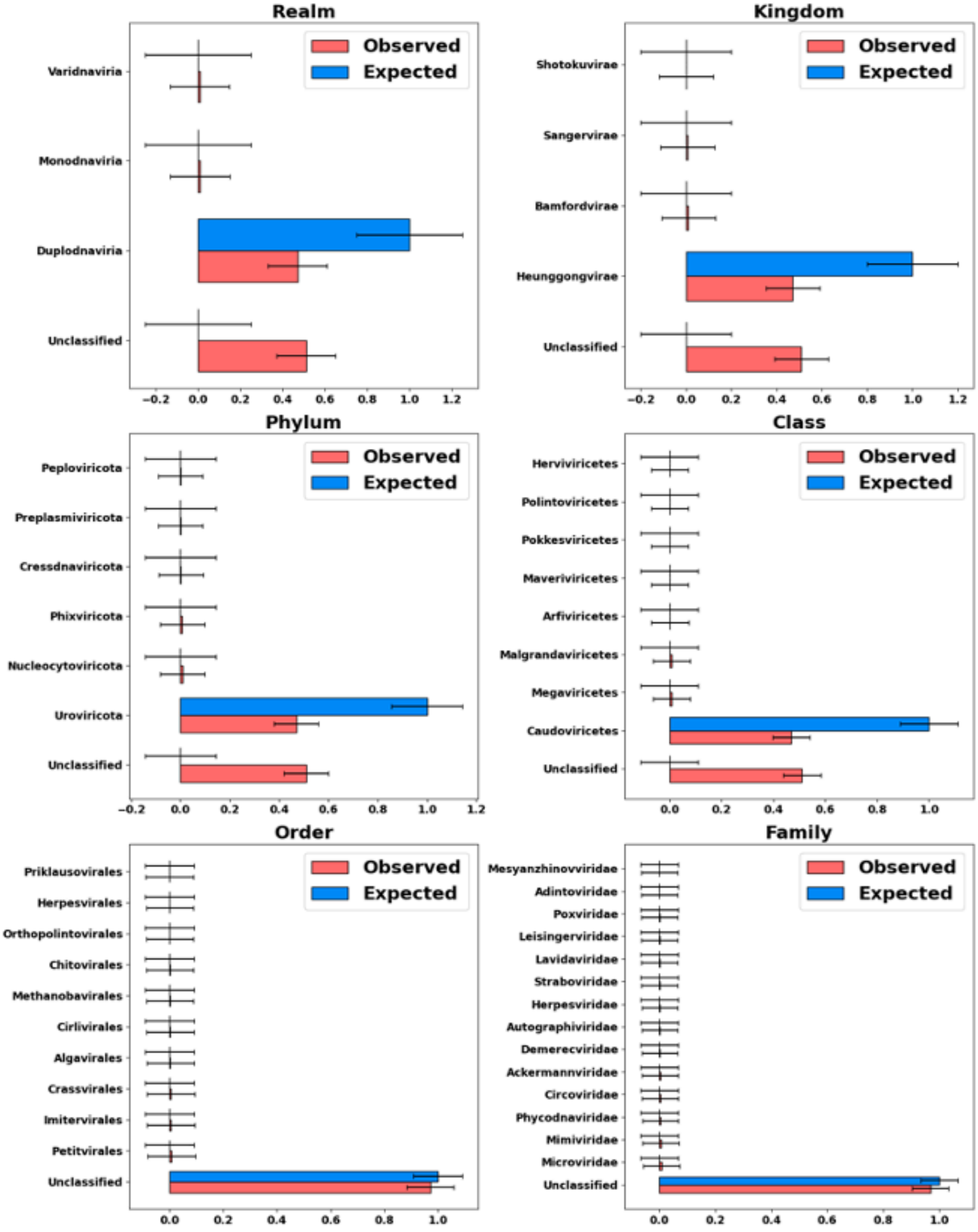


**Supplementary figure 22.** Comparison between IMG/VR counts (expected) and Prophage-DB counts (observed) for archaeal phages in host-associated environments. Random sampling with replacement was used (bootstrap value of 10,000).

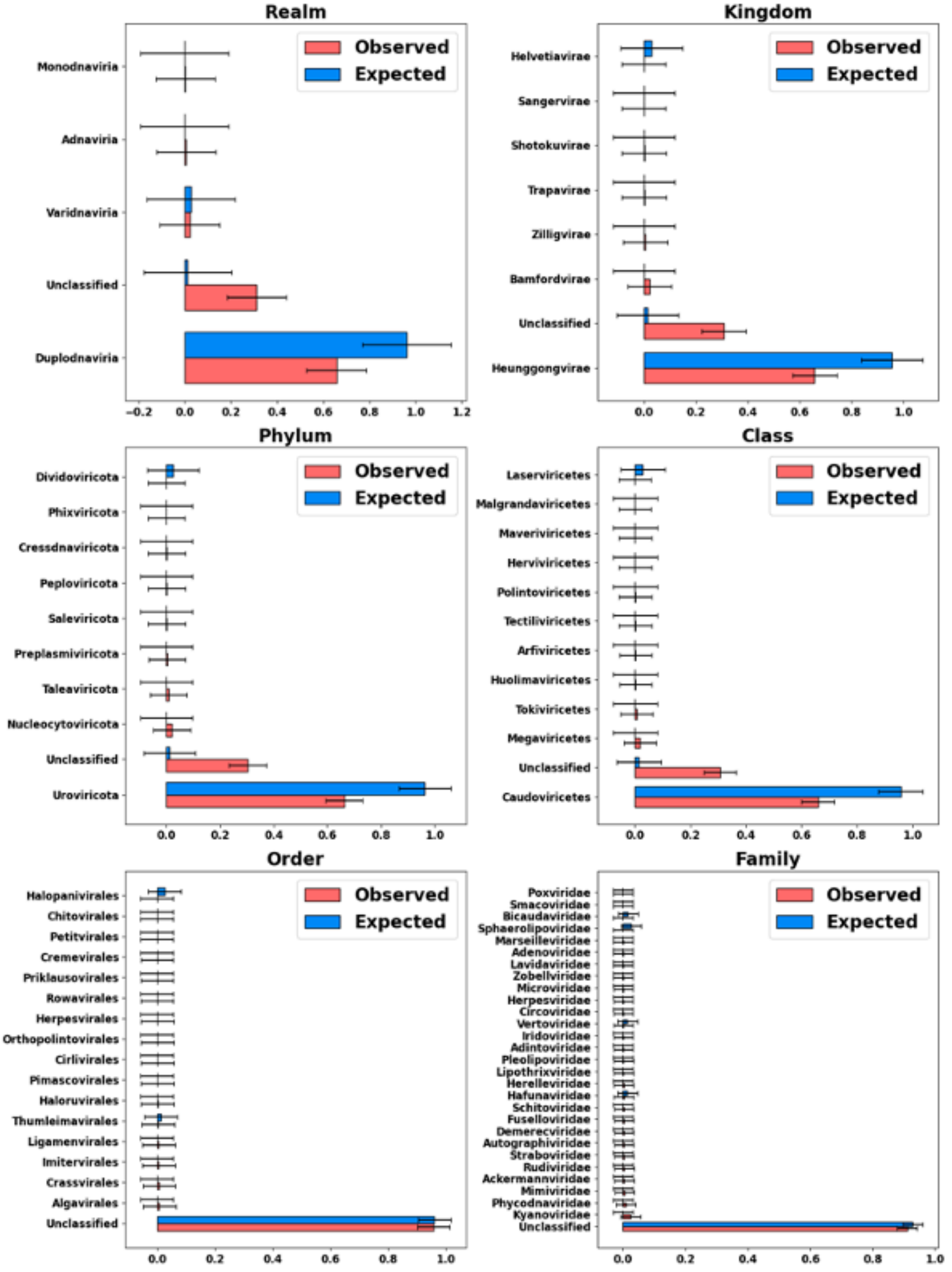


**Supplementary figure 23.** Comparison between IMG/VR counts (expected) and Prophage-DB counts (observed) for archaeal phages in marine environments. Random sampling with replacement was used (bootstrap value of 10,000).

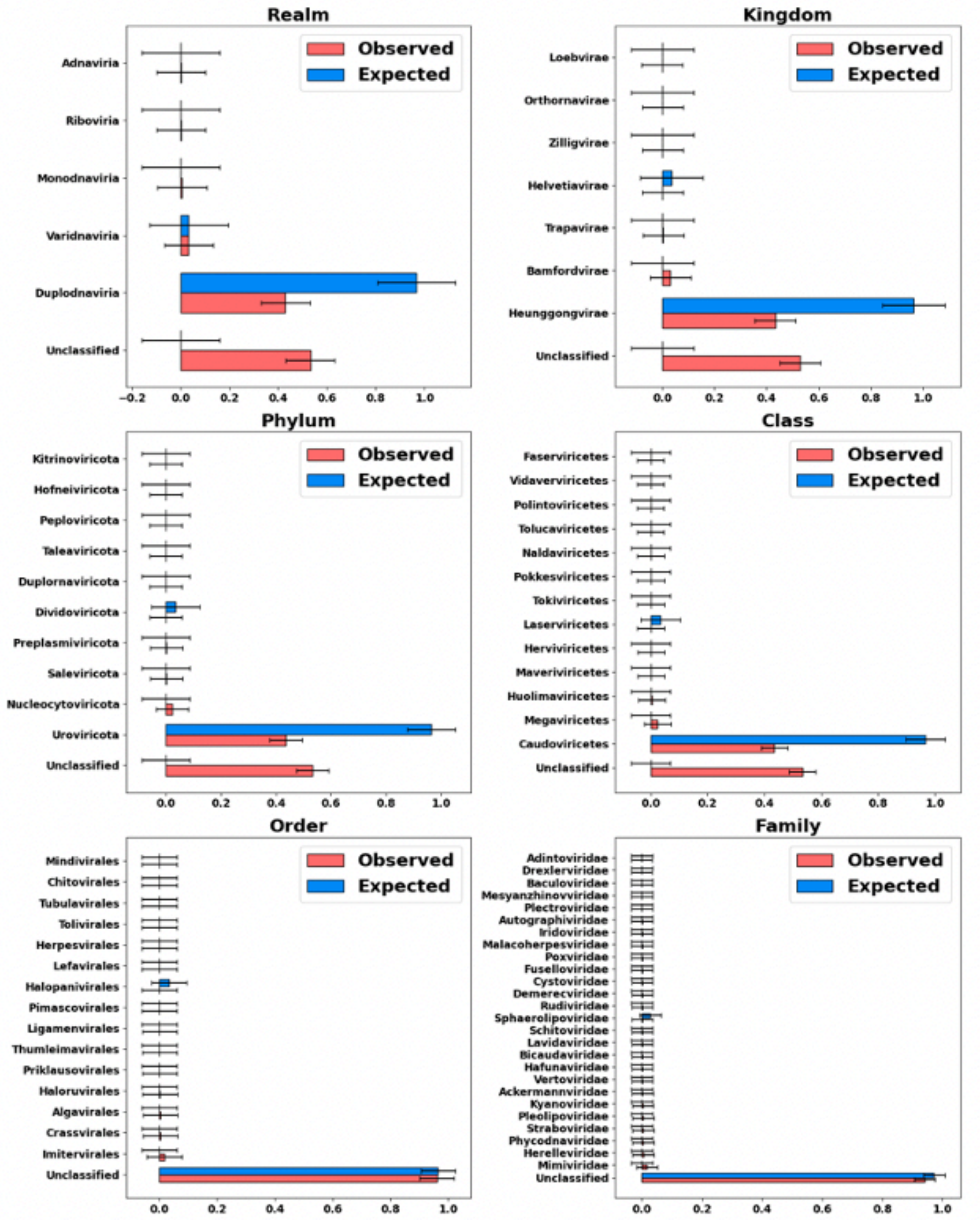


**Supplementary figure 24.** Comparison between IMG/VR counts (expected) and Prophage-DB counts (observed) for archaeal phages in terrestrial environments. Random sampling with replacement was used (bootstrap value of 10,000).
